# Deep sequencing of a large family of isogenic mice enables complex variants discovery and accurate phenotype mapping

**DOI:** 10.1101/2022.04.21.489063

**Authors:** Flavia Villani, Thomas Sasani, Mikhail Maksimov, M. Hakan Gunturkun, Nichole Ma, Yu-yu Ren, Daphna Rothschild, Hao Chen, Lu Lu, Beth Dumont, Kelley Harris, Melissa Gymrek, Vincenza Colonna, Jonathan K. Pritchard, Abraham A. Palmer, Robert W. Williams, David G. Ashbrook

## Abstract

The BXD family of recombinant inbred mice were developed by crossing and inbreeding progeny of C57BL/6J and DBA/2J strains. This family is the largest and most extensively phenotyped mammalian experimental genetic resource. Although used in genetics for 52 years, we do not yet have comprehensive data on DNA variants segregating in the BXDs. Using linked-read whole-genome sequencing, we sequenced 152 members of the family at about 40X coverage and quantified most variants. We identified 6.25 million polymorphism segregating at a near-optimal minor allele frequency of 0.42. We also defined two other major variants: strain-specific *de novo* singleton mutations and epoch-specific *de novo* polymorphism shared among subfamilies of BXDs. We quantified per-generation mutation rates of *de novo* variants and demonstrate how founder-derived, strain-specific, and epoch-specific variants can be analyzed jointly to model genome-phenome causality. This integration enables forward and reverse genetics at scale, rapid production of any of more than 10,000 diallel F1 hybrid progeny to test predictions across diverse environments or treatments. Combined with five decades of phenome data, the BXD family and F1 hybrids are a major resource for systems genetics and experimental precision medicine.

## Background

Understanding how genetic variation shapes biological traits, especially disease traits, remains one of the fundamental challenges in biology. This challenge lies at the heart of both evolutionary biology and precision medicine, yet our ability to connect genetic differences to phenotypic outcomes has been limited by the complexity of natural genetic variation. Deep whole-genome sequencing of genetic reference populations has become essential for understanding complex trait architecture and advancing precision medicine. Here, we report the comprehensive sequencing of the BXD family—the largest and most extensively phenotyped experimental genetic resource in mammals. Through linked-read sequencing of 152 BXD strains at ∼40X coverage, we have created an unprecedented catalog of genetic variants, from single nucleotide changes to complex structural variations. This resource, combined with five decades of accumulated phenotype data, enables us to move beyond traditional genetic mapping to precise variant-level predictions of phenotypic outcomes. Our work demonstrates how deep sequencing of a well-characterized genetic reference population can bridge the gap between genotype and phenotype, providing a powerful platform for experimental precision medicine.

Recombinant inbred (RI) strains are inbred strains that are produced by crossing and then inbreeding two or more inbred strains. In addition to mice [1–5], RI strains have been generated for numerous model systems, including yeast[6], plants[7,8], flies[9,10], nematodes[11], fish[12], and other rodents[13]. The BXDs RI strains are the largest mammalian RI panel and one of the oldest family of RI strains.

The first BXD strains were created in the early 1970s by crossing female C57BL/6J (B6) and male DBA/2J (D2) inbred mice[14]. The F2 progeny and all subsequent generations were then sibling mated to establish new “recombined” inbred lines (**Figure S1**). Each RI strain has a unique and stable genome that is a linear mosaic of ancestral haplotypes, inherited from *B6* and *D2* parents[15] (**Figure S1**). The first set of 26 BXDs were used mainly for mapping Mendelian loci[16,17], but as the family has grown—now to ∼152 members, they have found new uses, including mapping complex traits (phenotype-to-genotype), reverse genetics using phenome-wide association (genotype-to-phenotype), understanding gene-by-environment and gene-by-sex interactions[14,17,18], and causal modeling[19].

Two factors set the BXD family apart from other vertebrate genetic reference panels. First, the number of family members is large enough for well powered and precise genetic analyses[14]. There are now 120 strains available directly from The Jackson Laboratory (**Table S1**) with an additional 24 available from our laboratory at the University of Tennessee Health Sciences University. Second, the BXD has an extraordinarily rich multiomic phenome that has accumulated over more than 50 years. Virtually all of these “polyphenome” data are available from our large and FAIR-compliant[20] web service (GeneNetwork.org). This dense and well-integrated phenome consists of over 10,000 classical phenotypes[21] and well over 100 molecular and expression QTL-type data sets, with the ability to replicate deeply and to extend across many treatments and environments[22,23].

Although BXD progeny are all descended from the same B6 and D2 founders, because they have been created over several epochs, that are separated by several decades, the B6 and D2 parental strains have inevitably accumulated *de novo* mutations such that each epoch used slightly different B6 and D2 founders (**Table S1**). Furthermore, some of these epochs were produced using F_2_ intercrosses, while others used advanced-intercross progeny (**Figure S1A and S1B**). This has resulted in a hierarchical family genetic architecture that is not easily captured by genotyping arrays, but which is ideal for a family pangenome assembly — once all BXDs have been sequenced.

Since the first draft of the mouse genome in 2002[24,25], there have been important insights into the biology of the genome, the creation of new mouse models by genetic engineering[26], whole-genome screens, and the characterization of genomic diversity across mouse strains[27,28]. On a parallel course, stable mouse reference populations, such as the BXDs, have continued to be used for forward genetics, and with the sequencing of their parental strains, have also been used for systematic reverse genetics (also referred to as phenome-wide association studies)[9,29,30].

While inbred strains [24,27,28] have been sequenced previously [29,30], large, fully isogenic strain panels accompanied by rich, open phenome data have yet to be deeply sequenced. In the present study we have performed deep, linked-read sequencing of all the BXD family (mean read depth of 38.6X), enabling us to characterize the full complement of genetic variation within each BXD strain and epoch, including SNPs, small indels, short tandem repeats (STRs) and large structural variants (SVs > 1000 bp).

We also provide new, updated, and curated genotypes for a total of 198 BXD family members—from BXD1 through to BXD220, including the parental strains C57BL/6J and DBA/2J, 44 strains that are now extinct but for which phenotype data have already been acquired (**Table S1**), and whole-genome sequencing data for 152 of the extant strains. Deep sequencing reveals private variants unique to individual BXD strains and shared variants within BXD epochs, enabling precise mutation rate analysis and even mapping of mutation-controlling loci [31,32]. BXD progeny with extreme phenotypes can be caused by polygenic interactions or by private variants that likely occurred *de novo* during or after inbreeding was completed; for example, blindness in the BXD24-Cep90 line[26,28]. Whereas private mutations are a cause for concern when using RI lines for mapping, both epoch-specific and private variants, once identified, become valuable tools that bridge classic mutagenesis and quantitative genetic approaches. The result is an unprecedented compendium of variants within a single family, including many that are private to single strains or epochs. Combined with available phenome data, this resource enables us to transition from genetic dissection to genetic prediction and synthesis of genotype-to-phenotype relations.

## Results

### High resolution genetic analysis of the BXD mouse population using linked-read sequencing

We carried out linked-read sequencing from 10x Genomics on 152 extant BXD strains, plus their C57BL/6J and DBA/2J parents (**Figure S1**). Sequencing produced a mean read depth of 38.6X, a mean molecule length of 44.5 kb, and a mean 97% of reads mapped (**Figure S1C, Table S1**). The mean PCR duplication rate was 5.2%, indicating good library complexity. Coverage analysis showed that approximately 80% of the genome was covered at 20X or higher depth, with coverage declining more gradually at higher thresholds as shown in the coverage distribution curves (**Figure S1C).** Variants can be broadly separated into segregating variants - that is variants that are inherited from the C57BL/6J and DBA/2J parents of the population and segregate in the BXD family, and *de novo* variants - variants that have occurred in a particular BXD strain or group of strains during production.

### SNPs, indels and large structural variants

Using the GATK pipeline[33,34] and a set of true-positive variants in which the DBA/2J differ from the reference C57BL/6J, 5,891,472 SNPs and 696,596 indels have a minor allele frequency (MAF) greater than 0.2 and so we assumed to be inherited in the BXD family from the parental strains (**Figure S2B**). We clearly saw genomic regions of high diversity between the two parental strains (and therefore in the BXD family), as well as regions that were close to identical-by-descent between the parental strains (and therefore with few segregating variants in the BXD family; **Figure S2A**). The allele frequency spectrum reveals a distinct pattern typical of inbred strains, with most variants showing either very low frequencies, or a peak at 0.5, reflecting their origin from the parental genomes (**Figure 1A**). The size distribution of indels shows a symmetric pattern centered around zero, with a predominance of small insertions and deletions (**Figure 1B**).

**Figure 1:**
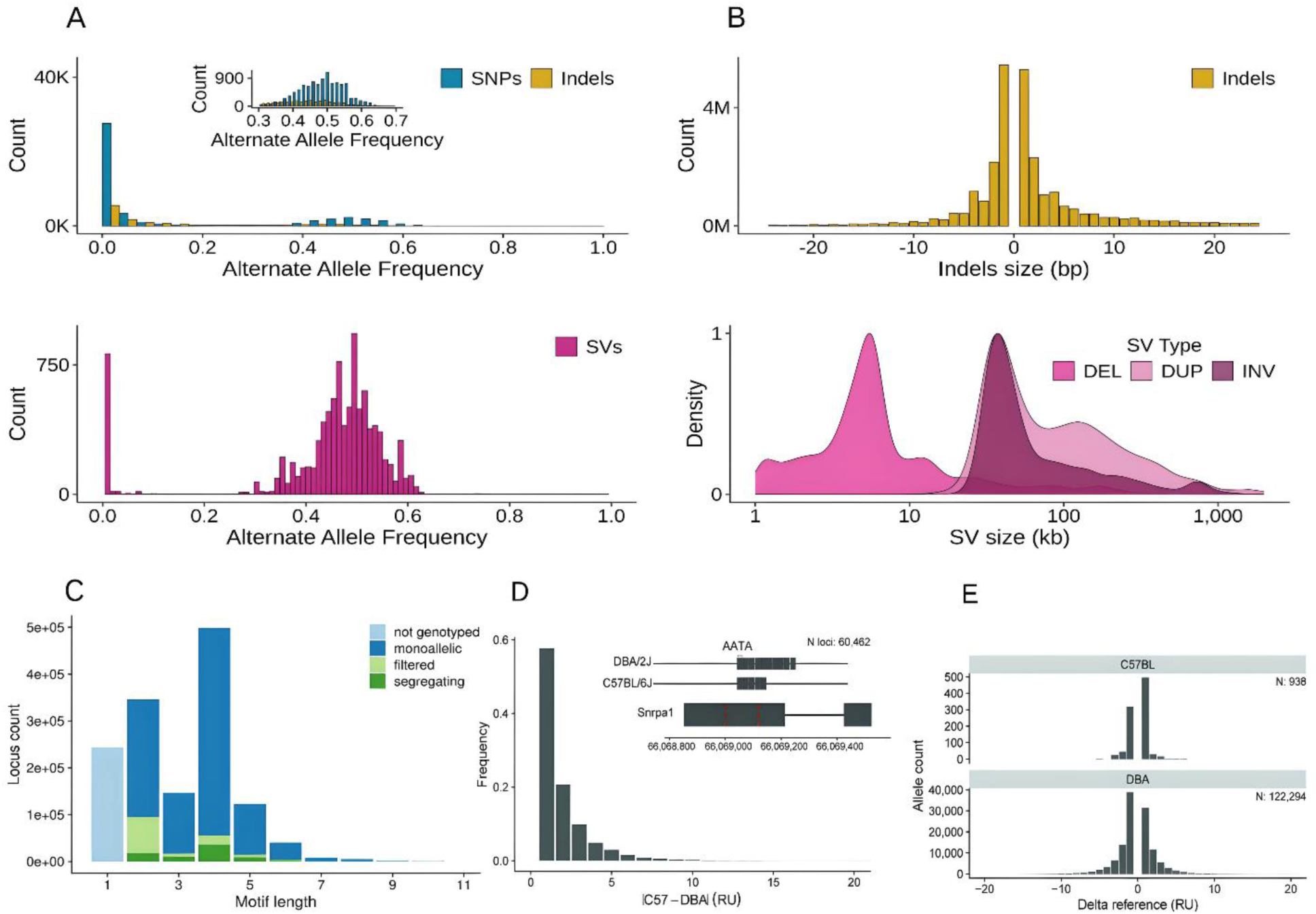
SNPs, indels and short tandem repeats (STRs) in the BXD family. **(A)** Allele frequency spectrum for three classes of variants: SNPs (blue), indels (gold) and structural variants (SVs, purple). Both SNPs and indels show a peak at low frequencies and a smaller peak at 0.5, while SVs display a unimodal distribution centered at 0.5 alternate allele frequency. **(B)** Size distribution of indels (gold) showing a symmetric pattern centered at zero, with most variants being small insertions or deletions (+/- 1-5 base pairs) and size distribution of structural variants by type: deletions (DEL), duplications (DUP), and inversions (INV), shown as density plots on a log scale (1kb to >1000kb). Deletions show a peak around 10kb, duplications and inversions a peak near 100kb. **(C)** Description of STR reference panel. Bar height indicates abundance of STRs of a given motif length. Homopolymers (light blue) were omitted from genotyping and monoallelic loci (dark blue) were omitted from downstream analyses. Calls not passing quality control metrics (light green) were filtered out to generate a final call set of segregating STR loci (dark green). **(D)** Distribution of differences in STR repeat counts between BXD founder strains and inset example locus with an AATA motif, overlapping an exonic region of a gene is shown as an example of a repeat where founder strains differ **(E)** Distribution of mutation size for STR loci relative to size of repeat in the reference for both parental strains—C57BL/6J and DBA/2J.

We used SVJAM[35] to jointly genotype large structural variants. We analyzed 31,454 candidate SVs suggested by the LongRanger pipeline[36] and called 4,153 SVs (1,968 deletions, 1,106 duplications and 1,079 inversions; **Figure S2C**). The allele frequency spectrum of SVs shows a unimodal distribution centered around 0.5 (**Figure 1A**) with SV size ranged from 1002 to 1,991,907 bp (mean 76,453 bp, median was 34,424 bp; **Figure 1C**). Notably, we found an SV hotspot with 272 SVs on chromosome 13. Of these, 128 were located between 64–67 Mb, and 56 were located between 67–70 Mb. This is a region with a high density of annotated segmental duplications, likely facilitating high rates of homology based structural variation (**Figure S2D**).

### Short tandem repeats (STRs)

To examine STRs, we built a genome-wide reference set of STRs with repeat units of 2- 20 bp in the GRCm38 (mm10) reference genome (**Figure 1D**)[37,38]. We then used GangSTR[37] to infer the repeat copy number at each STR in the reference in each BXD strain (Methods). Genotypes showed high call quality across repeat classes, with decreased quality at dinucleotide STRs (**Figure S3A**) which are more prone to errors introduced by PCR[39] in the 10x workflow. The majority of STRs in our reference panel were mono-allelic (69.3%) in the BXD and thus were excluded from downstream analyses. Another 8% of loci were filtered out either because they overlapped known genomic duplications or had an insufficient call rate across the family.

We used the resulting genotypes to characterize the extent of STR variation. As expected, *D2* showed increased variation relative to the reference assembly (**Figure 1E**). Considering only homozygous calls, the parental strains varied at 60,462 (78.8%) of polymorphic STRs (**Figure 1F**) with a mean difference in repeat length of 1.9 units. The remaining polymorphic STRs not found in the parents were predicted to be *de novo*, and are discussed below. While we found some larger repeat expansions and contractions of up to 10 repeat units, the majority had a difference of 1–2 units. Contractions relative to the reference allele were more abundant in D2 (55%/45%).

### Infinite marker map

A smaller set of 5,271,335 autosomal and 37,830 sex chromosome SNPs called with high confidence were used as the basis of an ‘infinite marker map’ that effectively defines every recombination point to the interval between the two nearest informative variants. This allowed us to identify almost every haplotype structure for each genome, and to identify the first and last variant in each haplotype block. We combined this with previous genotype-array data for extinct BXDs (**Figure S4A**). The total number of recombination per BXD ranged from 26 to 125 (**Figure S4B**). As expected, epochs derived from F2 crosses (epochs 1, 2, 4 and 6) have approximately half the average number of recombination as epochs derived from advanced intercrosses (epochs 3 and 5; **Figure S4B**).

The large numbers of recombinations in the BXDs, combined with the precise localization of each cross-over, increases the precision of QTL mapping. We reanalyzed the BXD phenome to demonstrate the advantages of these new genotype maps. For example, a phenotype that previously did not have a significant QTL (water intake of 13- week old females; GN BXD_12889), now has a significant and well-defined QTL on Chr 9 **Figure 2A**), encompassing only 5 coding genes. One of these genes, *Sorl1*, has a highly significant *cis*-eQTL in kidney. Although only 10 strains overlap between the gene expression dataset and the water intake, *Sorl1* expression and water intake are well correlated (Pearson *r*^2^=0.61) across those strains. Finally, variants in *SORL1* are associated with water intake in human PheWAS (*p*=0.00482; G/T, rs3781832, INI1528, https://biobankengine.stanford.edu/RIVAS_HG19/gene/SORL1). This demonstrates that the updated BXD genotypes allow *in silico* identification of novel gene loci with homologous function in mouse and human.

**Figure 2.**
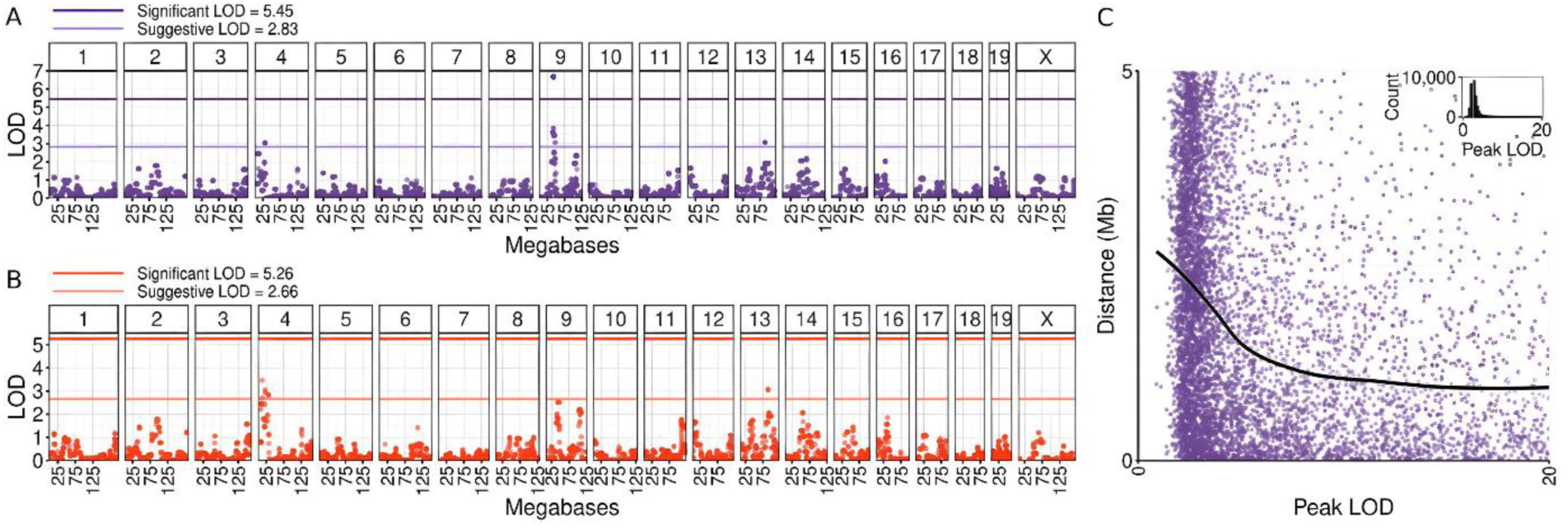
QTL mapping of water intake of 13-week old females (BXD_12889) using our new genotypes (A) and the old genotypes. **(B)**. The higher horizontal lines show genome-wide significance, whereas the lower horizontal lines show the suggestive threshold. Note that the QTL on Chr 9 is not even suggestive in B, but is highly significant in A. (**C) cis-eQTL accuracy in BXD mice using our new genotypes.** Each point represents the distance from the transcription start site of a gene (TSS) and the peak of a cis-eQTL for that gene, for LOD values between 3.5 (nominally significant) and 20 (highly significant). The black line is the best fit, showing the mean distance between the TSS and peak LOD. The inset shows a histogram of the number of cis-eQTLs with LODs between 0 and 20.

To estimate the overall precision of QTL localization, we used the largest available BXD transcriptome studies, containing at least 70 strains, encompassing 81 total distinct strains. Accuracy and precision were defined as the distance between the probe site and the peak QTL linkage value for all *cis*-eQTLs within 5Mb of the probe site. The large numbers of recombination in the BXDs, combined with the precise localization of each cross-over, increases the precision of the mapping (**Figure 2B**). Even for nominally significant loci (LOD 3.5), the median distance is 0.91 Mb, with the distance decreasing to 0.83 Mb at LOD > 4 and 0.74 Mb at LOD > 5. For these QTLs, the mean LOD is 8.94 and median LOD is 4.61, exceeding typical significance thresholds. This demonstrates that the distance between the peak of the QTL and the true causal variant will often be less than 1 Mb when using this new marker map. In summary, the new set of markers improves our ability to understand haplotype structures and more accurately identify recombination points in the BXD genomes. This feature greatly enhances QTL mapping accuracy and resolution, providing a valuable tool for both future genetic research in the BXDs, and the ability to reanalyze the phenome within the GeneNetwork integrated database.

### Accumulation and sharing of *de novo* mutations among BXD strains

#### Rates and spectra of new mutations - SNPs and indels

RI strains have always been used for mapping common parental variants, whereas rare or spontaneous fixed variants have been largely ignored. However, each member of the family is expected to carry its own unique set of homozygous mutations, which have arisen and been fixed over many generations of sibling matings (**Table S1**). To investigate rates and patterns of fixed *de novo* mutation among the BXD, we searched for all private homozygous singleton variants in each family member. These singletons are by definition absent from both DBA/2J and C57BL/6J parents and all other BXD progeny. We required these variants to be supported by at least 10 sequencing reads and to have a Phred-scaled genotype quality of at least 20. As expected, family members from earlier epochs have accumulated more homozygous singletons (**Figure 3A**). We additionally characterized the mutation spectrum of homozygous singletons, which describes the frequency of singletons corresponding to each of the six possible mutation types (C>A, A>G, etc.), in addition to C>T transitions at CG dinucleotides; overall, most singletons were C>T transitions (**Figure 3B**).

**Figure 3:**
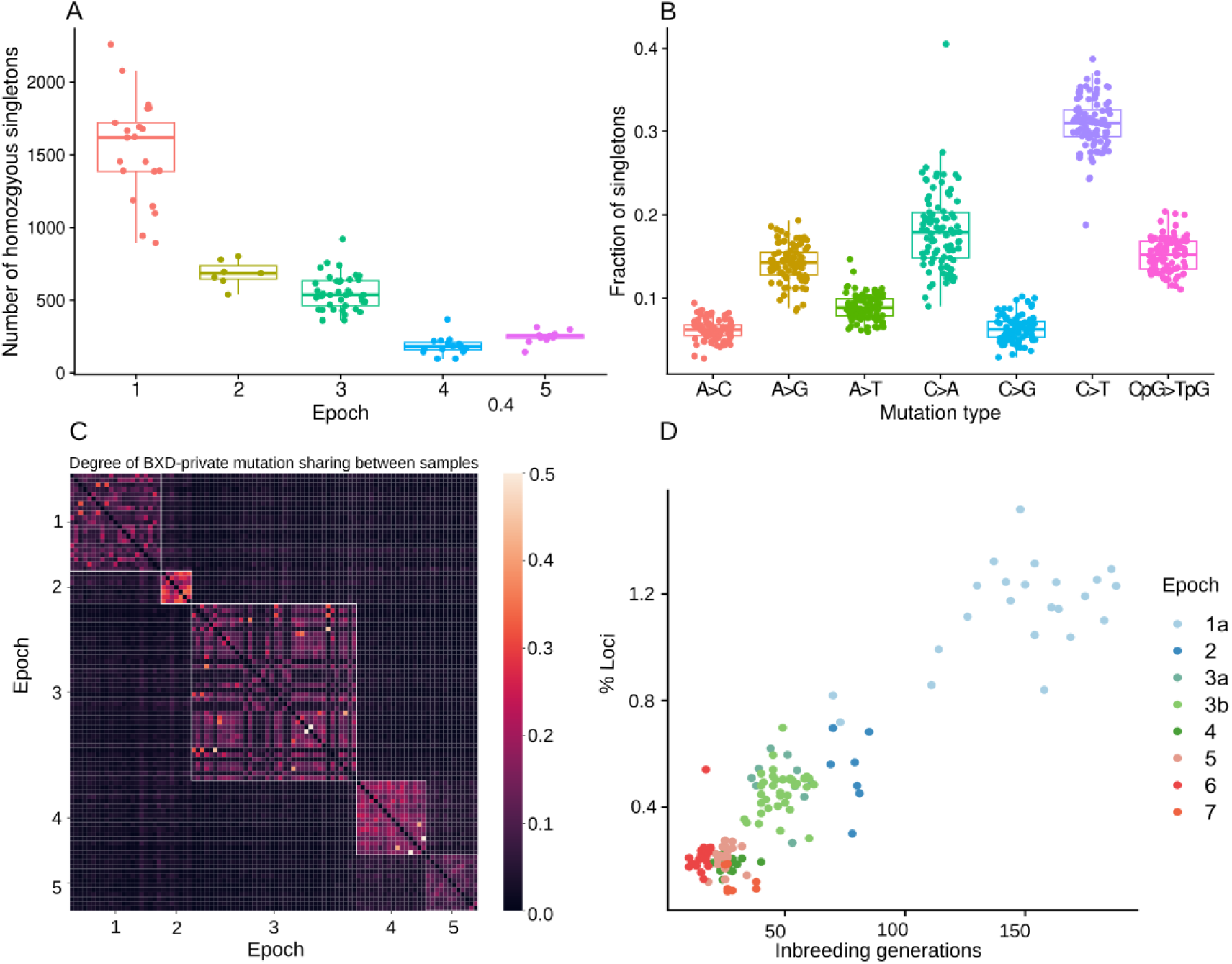
Germline mutation rates and spectra in the BXD family. **(A)** Counts of autosomal homozygous singleton SNPs in each BXD, stratified by epoch of origin. Average breeding generations are shown above each epoch. **(B)** Mutation spectrum of homozygous singleton SNPs across the BXDs, stratified by epoch of origin. Each homozygous singleton was classified according to its base substitution type, and C>T transitions were dichotomized by whether they occurred at CG dinucleotides. Mutations that were strand complements (e.g., C>T and G>A) were collapsed into a single mutation type. **(C)** Mutation sharing between BXD family members. We first identified mutations that were shared by 2 or more members of the BXD family, which likely represent recent *de novo* mutations private to a particular BXD lineage. For every pair of BXD family members, we then counted the number of mutations shared between the pair, and divided that number by the total count of mutations the two BXDs shared with any other BXDs; this fraction represents the degree of pairwise mutation sharing between the two BXD. Each epoch is outlined with a white box to highlight its self-similarity. **(D)** Correlation between number of inbreeding generations and percentage of accumulated non-parental STRs.

Some germline mutations might also arise earlier during the production of inbred strains and/or in particular founder individuals used for a particular epoch. Such mutations may be inherited by a subset of BXDs despite being absent from both the canonical C57BL/6J and DBA/2J parental strains. As each BXD epoch was initiated from a unique group of C57BL/6J and DBA/2J animals, we expected that these germline mutations would be shared more frequently by BXDs from the same epoch. On the other hand, *de novo* mutations shared by multiple BXDs from different epochs might represent recurrent *de novo* mutations or common genotyping errors. For each pair of BXDs, we calculated the number of sites at which they share mutations with each other, divided by the combined number of sites at which those two BXDs share mutations with any BXDs (in effect, a measure of Jaccard similarity between the sets of shared mutations in each BXD). Overall, BXDs tended to share more mutations with mice from the same epoch, demonstrating that some germline mutations are epoch specific, and were inherited by multiple RI strains (**Figure 3C**). Between epoch mutation sharing was observed between strains from epochs 4 and 5, potentially reflecting that these epochs were initiated from the same cryopreserved founders (from the JAX genetic stability program)[40]. Those founders may possess unique mutations absent from the mouse reference sequence[41].

### Rates and spectra of new mutations - STRs

STRs exhibit rapid mutation rates that are orders of magnitude higher than those of SNPs[42]. These mutations may occur at STRs that were fixed at a single allele in the founders or may occur at an STR that was polymorphic in the founders, leading to three or more alleles. Similar to other variant types, we expect recent *de novo* STR mutations to be heterozygous, whereas mutations arising in ancestors to present-day strains are more likely to be homozygous as a result of inbreeding.

We identified new mutations at STRs by searching for loci where BXD genotypes did not match either of the founder genotypes. The percentage of new variant loci per strain is highly correlated with the number of inbreeding generations (**Figure 3D**). Earlier epochs contain more than 1.2% *de novo* loci per strain, while more recently derived strains contain fewer.

### Sharing of de novo mutations across strains

Similarly to overall heterozygosity, the proportion of loci with one founder and one *de novo* allele increased in newer epochs, likely due to incomplete inbreeding (**Figure 3D**). However, these private heterozygous variants may also be enriched for genotyping errors.

To get a more reliable estimate of the number of loci in BXD strains which are not found in the parental strains, we considered only STRs where at least one strain had a homozygous non-parental genotype resulting in the identification of 18,135 unique STR loci harboring mutations (**Figure 4B**). As expected, the majority (54%) of non-parental mutations are singletons. The proportion of loci where we inferred a non-parental genotype for multiple strains drops off precipitously. As expected, many STR loci have three (the two parental alleles plus a mutation) or more alleles (**Figure S3B**). We validated a subset of observed homozygous new mutations using fragment analysis across 16 strains at 27 unique STR loci. Fragment analysis matched GangSTR at 364/374 of calls (97%) available in both datasets. Across 39 total *de novo* mutations tested, 35 (90%) were confirmed by fragment analysis. The majority of discordant calls were at a single locus (**Figure S3C**) and differed by a single repeat unit.

**Figure 4:**
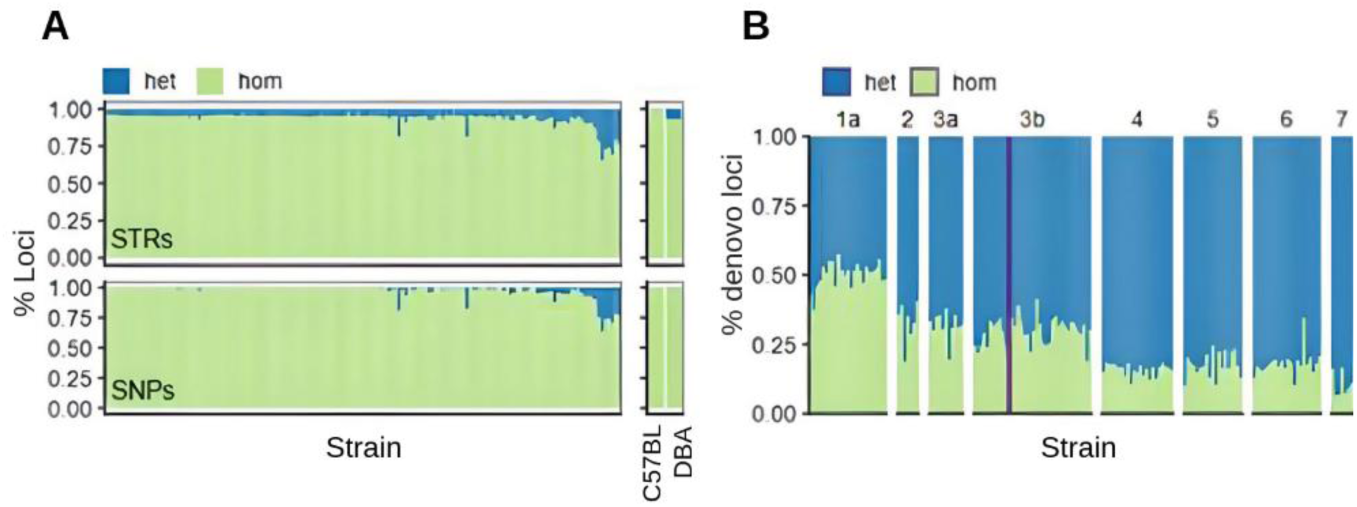
Short tandem repeat (STR) identification. **(A)** Comparison of the proportion of homozygous (green) and heterozygous (blue) STR calls for each RI strain (left panel) and founders (right panel) between SNPs and STRs. **(B)** Proportion of homozygous to heterozygous *de novo* STR calls for each RI strain.

Genotypes for the majority (92–96%) of variable STRs analyzed in both parents and BXD progeny were inferred to be homozygous for one of the two parental alleles (**Figure 4A**) and recapitulated the homozygous patchwork of inheritance blocks in agreement with the SNP results (**Figure 4B** and **Figure S4A**). Additionally, we observed an increase in heterozygous genotypes for family members from more recent epochs (**Figure 4A**), consistent with incomplete inbreeding.

### Identification of a novel QTL regulating short interspersed element (SINE) transposition rate

Approximately 40% of the mouse genome is composed of transposable elements, some of which remain active and contribute to genomic and functional diversity across laboratory strains[43,44]. SINE B2 elements are particularly abundant and active in mouse genomes, but the regulatory controls that limit their mobility to preserve genome integrity remain unknown [44]. To demonstrate our ability to find short interspersed elements (SINEs), we mapped B2 SINE counts using the newly generated marker map. We identified a significant QTL on Chr 5, peaking at 128.8 Mb, for the number of SINE elements. Although the QTL region was large (112.7 - 132 Mb) the peak was close to *Piwil1* (128.7 Mb), a gene that is known to promote the decay of a reporter mRNA containing a B1 or B2 SINE sequence[45].

#### Unraveling the role of private causal variants

Some private variants are already known in the BXD, especially those which cause extreme phenotypes. One example of this is the BXD24/TyJ-Cep290^rd16^/J strain. In 2004 a spontaneous mutation occurred in the BXD24/TyJ strain, causing a blindness phenotype[46–48]. Fortunately, frozen embryos from an earlier generation of the BXD24/TyJ strain were cryopreserved in 1988, allowing identification of an in-frame deletion in the causal gene, *Cep290*. The post-2004 strain was renamed BXD24/TyJ-Cep290^rd16^/J. Our study sequenced both of these strains, allowing us to confirm the variant and its position.

Whole-genome sequence enables rapid identification of causal variants using either a forward genetics approach (seeing variation in a phenotype, and finding the causal variant, as with BXD24/TyJ-Cep290^rd16^/J), or a reverse genetics approach (finding a genetic variant, and identifying phenotypes linked to it). Using variants identified in this paper, we provide examples of both approaches.

For example, we used a large liver proteome dataset (GN_AccesionId 887), and carried out Rosner Outlier Tests to identify outlier strains for each peptide. Among the results, we saw that BXD63 is an outlier for expression of three peptides (Q8BJY1_LEAPLEELR_2, Q8BJY1_VFTAIDQPWAQR_2, Q8BJY1_ELTGEDVLVR_2), all of which are part of the protein PSMD5 (proteasome 26S subunit, non-ATPase 5). We also saw that BXD63 had low expression of *Psmd5* in several liver transcriptome datasets. Examining our singleton variants in BXD63, we find a single variant, an insertion of a C (A > AC) at chr2:34860673, which Variant Effect Predictor (VEP) annotates as a regulatory region variant. We hypothesize that this insertion is the cause of low *Psmd5* expression and low levels of PSMD5-derived peptides observed in BXD63.

BXD29, for which we sequenced two ‘sister’ strains, is an excellent example of how our new sequence data can solve old questions. It has previously been discovered that the BXD29-*Tlr4^lps-2J^*/J mouse strain has a highly penetrant bilateral nodular subcortical heterotopia and partial callosal agenesis, and that it is not caused by the *Tlr4* variant[49,50]. In our sequencing of the BXD, we have identified only 975 variants that are present in the BXD29-*Tlr4^lps-2J^*/J mouse strain that are not shared by the BXD29/TyJ strain - a strain derived from cryopreserved bankstock of F39 from 1979. From breeding records, we know that a spontaneous mutation arose in the BXD29-*Tlr4^lps-2J^*/J between 1998 and 2004[49,50]. Given the large effect of the variant, we hypothesize that it is protein-coding, leaving only 30 of the 975 variants as candidates. These include a large duplication (chr11:74,689,025-75,132,234) affecting seven genes (*Mettl16*, *Mnt*, *Pafah1b1*, *Sgsm2*, *Smg6*, *Srr*, *Tsr1*). Importantly, in the human genome, duplication of this region, especially *Pafah1b1,* causes cortical defects[51,52], including subcortical heterotopia^[^53,54^]^. A F_2_ cross between the BXD29/TyJ and BXD29-*Tlr4^lps-2J^*/J (sometimes termed a reduced complexity cross[55]) could be used to formally confirm this.

#### Pangenomic analysis identifies complex variants and is informative about strain-specific haplotypes

All the above work has used the classic approach of aligning to a reference genome. However, pangenomes[56–59] have key advantages, allowing all known variants to be included as part of the alignment process. As such, we built the BXD pangenome of Chr 19. This is expected to improve mapping of sequencing data (e.g. whole genome sequencing, RNA-seq and methylation-seq), and allow us to discover new variants. The BXD pangenome for Chr 19 consisted of 5.5M nodes and 8.6M edges (total length 264,380,676 bp with 82,695 paths, **Figure 5A**), starting from haploid assemblies of 10X reads. Four strains were excluded for poor assembly quality, leaving 148 strains in the pangenome.

**Figure 5.**
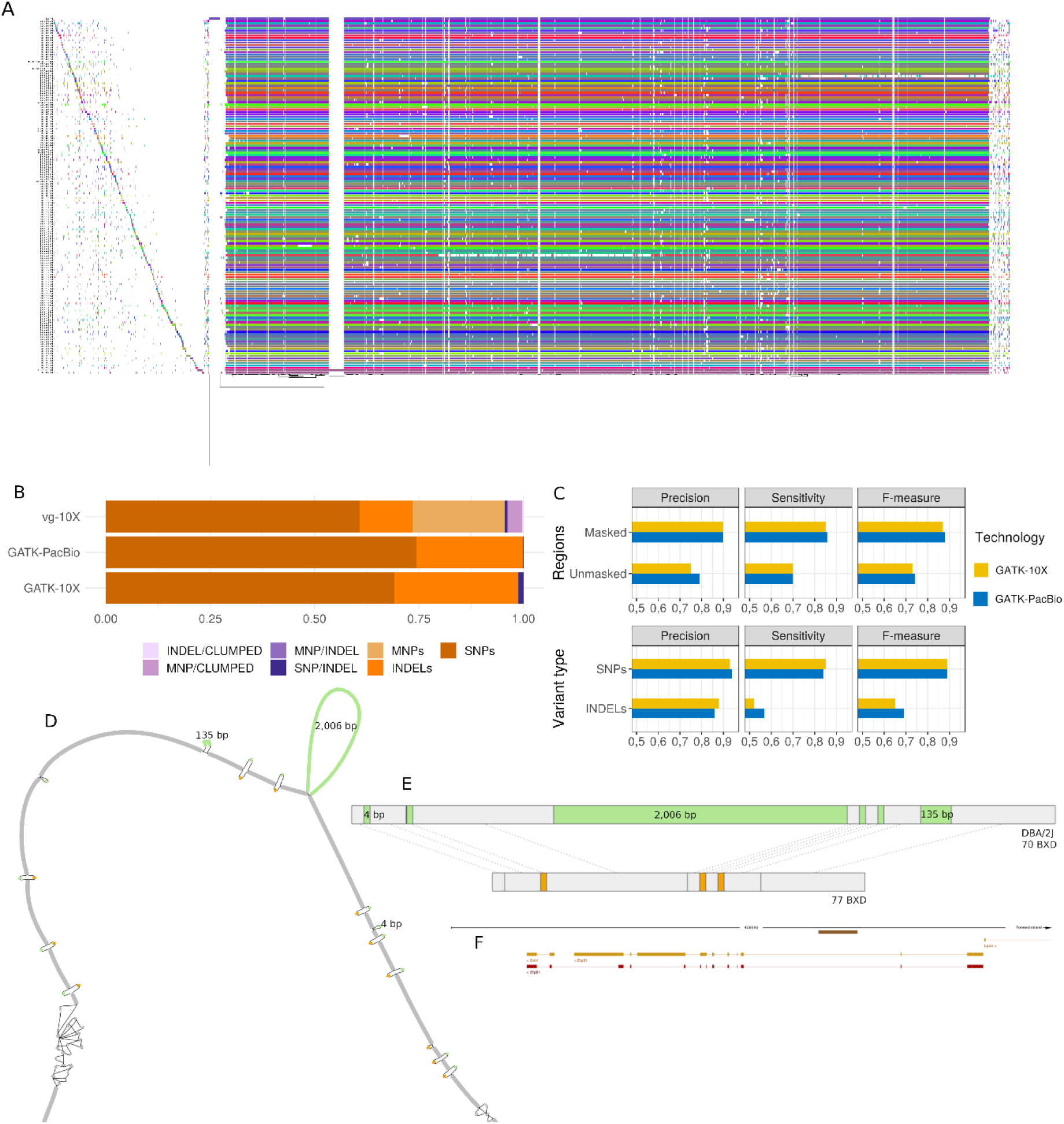
Pangenome of the BXD family. (A) *odgi-viz* linear visualization of the pangenome of Chr 19. Each line represents a haplotype. Line interruptions (white) are insertions in one or more strains, therefore deletions in the others (vertical white stripes). The left side is the centromere. In these two regions sequences are fragmented. **(B) Types of microvariants** (up to 50bp) at variable sites in the pangenome call set (vg-10X), and the two truth sets (GATK-PacBio and GATK-10X). Simple variants are in shades of orange, complex in shades of purple. *vg* call set includes the higher fraction of complex variants**. (C) Evaluation of *vg* call set in DBA/2J** using two truth sets sequenced with different technologies and called with GATK. Evaluation by variant type is within masked regions. **(D) Extract of the pangenome from the *Zfp91* gene** showing a 2,006 bp insertion found in DBA/2J and 48% of the BXD strains (green nodes in the graph). The insertion is in complete linkage with two other insertions of 4 bp and 135 bp in a region spanning 2.8 kbp. **(E) Strain-specific haplotypes** (gray segments are not in scale) **(F) Pangenomic extract in the gene context** is represented here by the brown segment above the second intron of the gene. The region on display corresponds to Ensembl *M.musculus* version 102.38 (GRCm38.p6) chr19:12,758,303-12,800,960.

We first focused on genetic variants up to 50bp (microvariants; SNPs and small indels). In DBA/2J we identified 182,643 variable sites. Of these, 178,557 are simple variants (SNPs and indels), and 4,086 are complex variants (allelic variants that overlap but do not cover the same range; **Table S2),** many of which were not detected by GATK in our reference based variant calling (**Figure 5B**). This shows that the pangenome enables calling of complex variants such as clumped multiple nucleotide polymorphisms (MNPs) and MNP/INDEL, variants not detected by traditional genomics methods(**Table S2**). The Ts/Tv ratio in the pangenomic calls is 2.08, which is slightly lower than figures from GATK (2.15 and 2.09 for 10X and PacBio data, respectively), reflecting the enrichment of complex variant calls in the pangenomic set. Within masked regions (i.e. regions that do not contain SINE, ALUs, LINE, LTR, and other DNA repeats), precision and sensitivity of vg[60] calls are 90% and 85-86%, respectively, when compared to GATK. For SNPs these figures rise to 93-94% and 84-85% (**Figure 5C**, **Table S3**), matching precision obtained from the comparison of the two truth sets (**Table S3**).

Considering variants >50bp, we identified 61,381 variable sites. One example of these is a 2kb insertion (**Figure 5D**) in the second intron of the E3 ubiquitin ligase *Zfp91* (**Figure 5E**). This insertion was described in the DBA/2J strain[28] and is found in 48% of our BXD mice in complete linkage with two other insertions of 4 bp and 135 bp on a haplotype spanning 2,789 base pairs (**Figure 5F**). Using the presence or absence of the SV as a phenotype in GeneNetwork, we correlated this against the BXD phenome. The second most significant correlation was with ‘Thymic T-cell proliferative unresponsiveness (anergy) to anti-CD3-induced proliferation’ (p = 1.095e-04, r = 0.719). The same association between *Zfp91* and T-cell proliferation has recently been made by others in gene knockout mice[61]. The *Zfp91* gene facilitates TCR-dependent autophagosome formation to sustain T reg cell metabolic programming and functional integrity. *Zfp91* deficiency attenuates the activation of autophagy and associated downstream pathways and impairs T reg cell homeostasis and function, rendering mice sensitive to colonic inflammation and inflammation-driven colon carcinogenesis[62]. Therefore, there is strong evidence that segregating variants in *Zfp91* in the BXD cause alterations in T-cell proliferation, and we expect this to result in clinically relevant phenotypes, e.g. relating to cancer and the immune system.

A similar analysis performed for pangenome of chromosome 13, contributed to the identification of SVs as part of a recent Genome Research paper[32]. Overall, while the ideal data for an effective pangenome construction is long-reads, we conclude that the pangenome produced by short linked-read can be informative in model organisms[63].

## Discussion

We report the sequencing of the entire BXD family, bringing deep, whole-genome, linked-read sequencing to this widely used and deeply phenotyped resource. We have identified SNPs, small indels, large structural variants, short tandem repeats and SINEs.

Using the known family structure of the BXD and the deep phenome that has been collected, we are able to link these variants to phenotypes. We show how these can be used to find new, molecular phenotypes, and to improve our ability to find causal genes for classical phenotypes.

The BXDs are well suited to studying gene-by-environmental interactions (GXE) and for experimental precision medicine[14]. For almost 50 years, they have been used to study the genetic and environmental factors that underlie a diverse collection of phenotypes, including environmental toxicant exposures, alcohol and drugs of abuse[64–68], infectious agents[69–71,71,72], diets[73–78], and stressors[79,80]. Beyond this, there are also extensive -omics data available for many BXD family members, including over 100 transcriptome datasets (e.g,[81,82]), as well as more recent miRNA[83,84], proteome[85–87], metabolome[82,87,88], epigenome[17,89], and metagenome[90,91] profiles. As each of these new data sets is added, it can be integrated with previous data, thereby multiplicatively increasing the usefulness of the whole phenome. We can easily identify strains that are outliers for any particular trait or molecular phenotype, and immediately have a short-list of candidate variants.

Although the BXD have been well characterized by array-based methods, allowing a long history of QTL mapping, this is the first time that the full breadth of variants have been available for the family. SINEs[92], CNVs[93], and large deletions[18] have been found in the family previously, but these required extensive work to get from locus to variant. With the whole catalogue of variants available this is now only an afternoons’ work.

Using deep genome sequencing data, we have also been able to investigate variation in mutation rates and spectra across the BXD family[25,32]. These called variants can become molecular phenotypes, which themselves can be mapped to causal loci. For example, we are able to identify genes causing specific mutation signatures[31,32], and a locus influencing the number of SINEs in the genome. We give a brief overview of some of these analyses here, but they are an interesting and important outcome in and of themselves, and will provide fuel for future research.

The BXD are an enduring resource: phenotypes collected in the 1970s can be mapped using genotypes identified by our study, leading to the identification of novel QTLs. By extension, phenotypes collected in 2025 will still be informative in 2075. This is made possible not only by the use of well-characterized inbred strains, but by the dedicated community who have donated their data to open access resources, particularly GeneNetwork[22,23,87–94]. In this way, phenotypes collected across decades, continents and environments can be coherently coanalyzed, providing new insight from legacy data[94].

Furthermore, by crossing two BXD strains any of 22,350 isogenic F1 hybrids can be generated – a massive diallel cross (DAX)[14]. Thus, there are well over 22,000 reproducible “clones” of F1 hybrids that can be generated from the current BXD families, each of which carries one chromosome from each of its BXD parents, and is therefore fully ‘*in silico*’ sequenced in advance. This will be a huge resource for the understanding of indirect genetic effects, gene-by-environment interactions, and epistasis. These can also be crossed to any other inbred strain, such as the Collaborative Cross population[1–5], or genetically engineered mice[95–97], expanding the amount of variation, and allowing for identification of modifier alleles. This has been excellently demonstrated with the AD-BXD[98–101], and others[102,103].

Given the deep and well recorded history, with over 180 generations of inbreeding in some strains, we have a unique resource. This project revitalizes this 50-year-old family, allowing many new analyses beyond the sample given here, and exponentially expands the utility of other data collected over many decades. This data not only allows the identification and genetic mapping of ‘molecular phenotypes’ (e.g. mutation spectra), but is fully interactable with the whole BXD phenome, and is available for all future users of the BXD family.

In the context of genomic research, it has become increasingly evident that relying on a single reference genome can introduce significant biases and limitations that impact various aspects of genetic analysis. These biases can have adverse effects on variant discovery, gene-disease association studies, and the overall accuracy of genetic analyses. One example of this issue is the reference genome for DBA/2J, which, unlike the well-established C57BL/6J reference, lacks the same level of quality and completeness. Utilizing a lower-quality reference assembly for DBA/2J can introduce its own set of errors and biases into genomic analyses.

To address these challenges and to advance the field of genomics, there has been a growing interest in transitioning from the use of single reference genomes to the adoption of pangenomes. Pangenomes represent a paradigm shift in genomic analysis, offering a more comprehensive and flexible approach. They encompass a broader spectrum of genetic diversity within a given species, capturing not only the common features found in most individuals but also the unique genomic variations that distinguish individuals from each other. In the near-future, sequencing technologies such as improved and cheaper long-read sequencing, offer the promise of constructing higher-quality assemblies and pangenomes, quickly and easily. These technologies provide a more comprehensive view of genomic diversity and reduce reliance on a single reference genome. This shift has the potential to enhance our ability to discover genetic variants, unravel complex gene-disease associations, and improve the precision and reliability of genetic analyses in the future.

## Conclusions

The deep sequencing of the BXD family at ∼40X coverage represents two major breakthroughs in systems genetics. First, we achieved unprecedented precision in mapping recombination breakpoints in the BXD, revealing exactly where and how genetic material exchanges occur across generations. Second, using linked-read technology, we established a comprehensive catalog of genetic variants that far surpasses previous efforts, capturing three distinct classes of variation: founder-derived polymorphisms, strain-specific *de novo* mutations, and epoch-specific variants. This technical achievement provides a level of detail that was previously invisible to traditional short-read sequencing, eliminating guesswork from variant identification.

By integrating this precise genetic data with five decades of accumulated phenotype data, we’ve created more than just a reference database - we’ve developed a powerful platform for experimental precision medicine. The resource enables researchers to systematically analyze over 22,000 possible F1 hybrid combinations, each fully characterized at the sequence level before any experiments begin. This capability, combined with the extensive phenome data available through GeneNetwork.org, changes how we approach genotype-phenotype relationships, allowing precise variant-level investigations across diverse environments, treatments, and genetic backgrounds. The result is a transformative tool for understanding how genetic variations shape the diverse phenotypes observed in the BXD family, setting a new standard for genetic reference populations.

## Acknowledgements

We thank Dr. Benjamin A. Taylor for initiating the BXD and for continued support and encouragement. We thank Drs. Gerald McClearn and Lisa Tarantino for 60 eighth-generation (G8) AI progeny from Pennsylvania State University that contributed to epoch 3, in addition to B6D2 AI progeny (G8 and G9) to make BXD160 through BXD186 contributed by Dr. Abraham Palmer. We thank Dr. Andrew Clark for his helpful discussion.

The UTHSC Center for Integrative and Translational Genomics (CITG) has supported production of the BXD colony at UTHSC and will continue to support this colony for the duration of the grant. The CITG also provides generous support for computer hardware and programming associated with GeneNetwork, and our Galaxy and UCSC Genome Browser instances. We thank the support of the UT Center for Integrative and Translational Genomics and funds from the UT-ORNL Governor’s Chair, NIDA grants P30DA044223, P50DA037844 and U01DA051234; NIAAA grants U01AA013499, U01AA016662, and U01AA014425. The BXD Resource at the Jackson Laboratory is supported by NIH P40OD011102 awarded to Dr. Cathleen M. Lutz. The work at Stanford University is supported by NHGRI grant R01HG008140. We thank the The University of Tennessee Infrastructure for Scientific Applications and Advanced Computing (ISAAC) for hosting raw data and compute power. The data sets generated and/or analyzed during the current study are available in the GeneNetwork repository, https://www.genenetwork.org/.

## Author contributions

Conceptualization: J.K.P., A.A.P., R.W.W.

Methodology: F.V., T.S., M.M., M.H.G., B.D., D.G.A.

Software: FV, T.S., M.M., M.H.G., H.C., B.D., D.G.A

Formal Analysis: F.V., T.S., M.M., M.H.G., N.M., R.Y., D.R., B.D., D.G.A

Investigation: F.V., T.S., M.M., M.H.G., N.M., B.D., D.G.A

Resources: L.L.

Data Curation: D.G.A., R.W.W.

Writing – Original Draft: D.G.A.

Writing – Review & Editing: F.V., T.S., M.M., M.H.G., F.V., H.C., V.C., B.D., K.H., D.G.A

Visualization: F.V., T.S., M.M., M.H.G., N.M., F.V., B.D., K.H., D.G.A

Supervision: H.C., V.C., B.D., K.H., J.K.P., A.A.P., R.W.W.

Project Administration: D.G.A., J.K.P., A.A.P., R.W.W.

Funding Acquisition: J.K.P., A.A.P., R.W.W.

## Methods

### Contact for Reagent and Resource Sharing

Further information and requests for reagents may be directed to, and will be fulfilled by, the corresponding author David Ashbrook.

### Data and code availability

Sequence data have been deposited at European Nucleotide Archive and the Sequence Read Archive and are publicly available as of the date of publication. under project PRJEB45429. This paper does not report original code. Any additional information required to reanalyze the data reported in this paper is available from the lead contact upon request.

### Experimental Model and Subject Details

*Tissue was taken from 154 males, mainly young adults. Full details of all individuals are in Table S1. All BXD strains are available under a standard material transfer agreement; the most important limitation being that they cannot be sold or distributed without approval of the Jackson Laboratory or UTHSC. Availability information on all BXD strains are in Table S1. Mice were euthanized using isoflurane, tissue was collected immediately, flash frozen with liquid nitrogen, and placed in the -80 freezer for later analysis*.

### Method details

#### Sequencing the BXD families

DNA was extracted from 50 to 80 mg of tissue. DNA extraction, library preparation and sequencing was carried out by HudsonAlpha. High molecular weight (HMW) genomic DNA (gDNA) was isolated using the Qiagen MagAttract kit. The Chromium Gel Bead and Library Kit (v2 HT kit, revision A; 10X Genomics, Pleasanton, CA, USA) and the Chromium instrument (10X Genomics) were used to prepare the libraries for sequencing. The barcoded libraries were sequenced on an Illumina HiSeq X10 system.

#### Quantification and Statistical Analysis

The phasing software LongRanger (v2.1.6)[36] was run to generate a phased call-set of single nucleotide variants (SNVs), insertion/deletions (indels), and structural variant discovery, against the mm10 reference genome.

Joint calling used GATK (v4.0.3.0)[104]. HaplotypeCaller was used to create joint caller gvcf files from bam files produced by LongRanger. Variant calls use family-wide information, increasing the likelihood of detecting segregating variants—either between the parental strains, or within the subfamily epochs.

Variant quality was calculated using *variant quality score recalibration* (VQSR) from GATK, the inputs are gVCFs files and the mm10 reference genome. A list of known variants was produced by finding those variants we detected in all three of the following independent resources: 1. our own new sequencing of DBA/2J; 2. Wang et al (2016)[18] sequencing of DBA/2J using both SOLiD and Illumina; and 3. the Sanger Mouse Genomes project[28] sequencing of DBA/2J using Illumina. The union of these three includes 3,972,727 SNPs, 404,349 deletions and 365,435 insertions. As expected, these variants segregate with a MAF close to 0.5. This set has been taken as a validated dataset. A training dataset was created using the mgp.v5.merged.snps_all.dbSNP142.vcf.gz (5/12/15) and mgp.v5.merged.indels.dbSNP142.normed.vcf.gz (4/30/15) files from ftp://ftp-mouse.sanger.ac.uk/current_snps.

#### Structural variant calling

We developed a joint calling method, SVJAM, that detects large structural variants (SV) from linked-read data across multiple samples. A detailed description of the algorithm is available from Gunturkun et al. 2021[35]. Briefly, SVJAM first collects candidate SV regions from individual samples reported by LongRanger[36], which is error prone. We then retrieve barcode-overlapping data for each candidate location from all samples using the Loupe application of the 10x Chromium Platform (**Figure 6A, 6B**), one image for each individual from a genomic location of interest. The intensity of pixels in these images represent the depth of barcode overlap for the corresponding genomic locations and are the primary data for our analysis.

**Figure 6:**
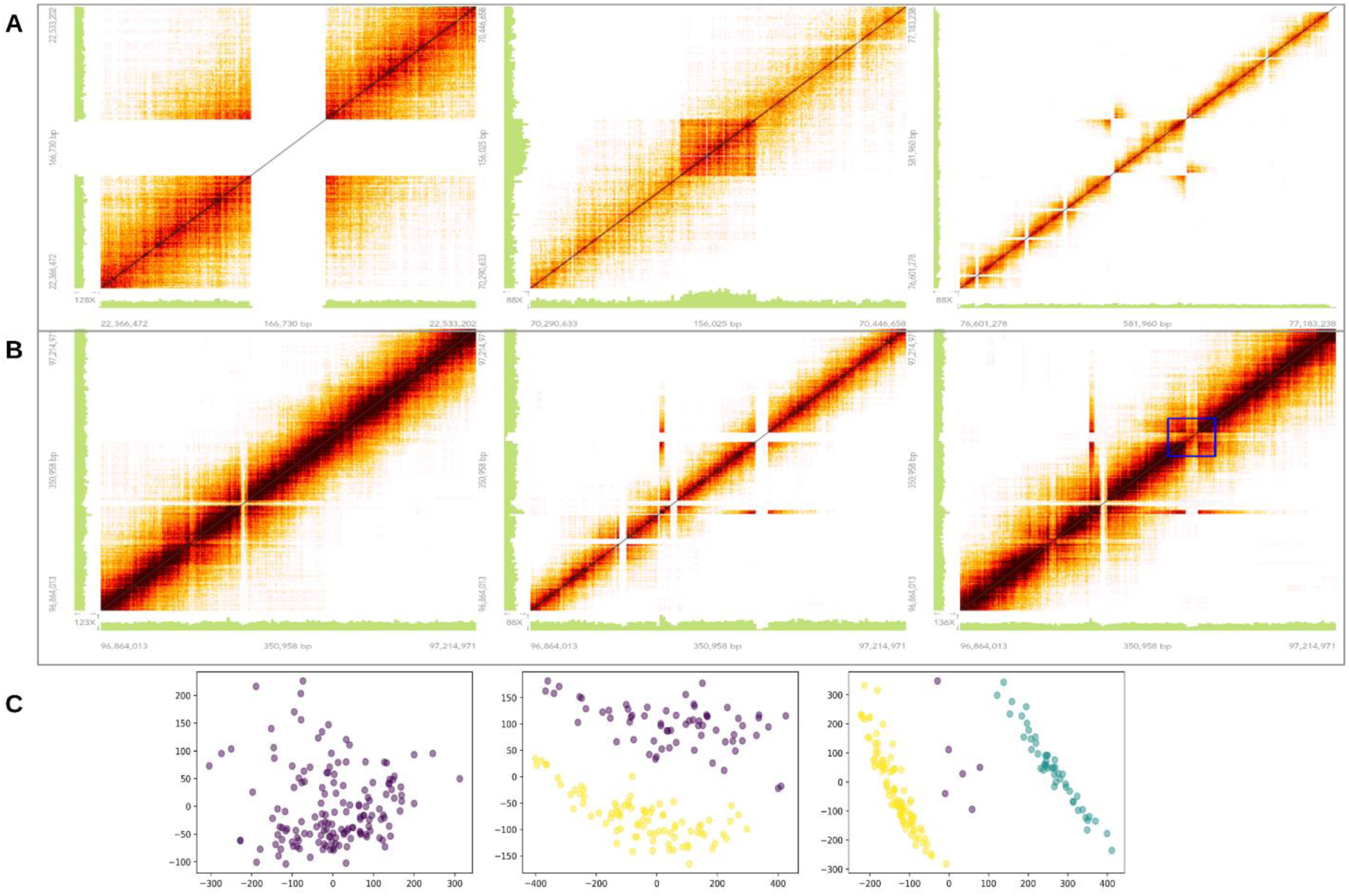
Joint calling of large SVs from linked-read data. **(A)** Example barcode overlapping images of deletion, duplication and inversion. **(B)** Examples of homozygous reference, homozygous alternative, and heterozygous variants.**(C)** Joint calling showing one, two or three genotypes. Samples (showing as dots) are projected onto a two dimensional space after principal component analysis. Colors represent genotype calls obtained using a clustering algorithm with a custom distance matrix.

We used slightly different processes for different types of SV. Conventional image processing techniques are used to detect deletions, which are represented by symmetric gaps with no barcode overlap along the diagonal of the image (**Figure 6A**, first image). The beginning and end of the deletion are determined according to the location of the gap on the x- and y-axis. Duplication (**Figure 6A**, second image) and inversion (**Figure 6A**, third image) are called using a convolutional neural network (CNN) with convolution, max pooling, dropout and flatten layers. This CNN is trained on a set of 12,000 images with validated labels (i.e. type of SV).

To further increase the accuracy of genotype calls, we designed a joint calling procedure that determines the presence of structural variants and the genotype of each individual. This procedure first converts each image into a matrix and then flattens it to a vector. The vectors of each sample are then combined in columns to form a matrix having approximately one million (pixels per image) times 152 (number of samples) dimensions for each candidate location.

The primary algorithm of the joint calling is a principal component (PC) analysis on the high dimensional matrices. Samples projected onto a two-dimensional space of the first two PCs display different patterns based on the number of genotypes: no distinctive pattern is shown when only one genotype is present, two distinctive clusters are shown when there are two genotypes. Because the BXD panel is inbred, when heterozygotic samples are present, they always contain a small number of samples and are located between the two homozygotic clusters (**Figure 6C**).

We applied hierarchical clustering to the data projected onto the 2D plane to call the genotypes. We designed a custom metric *d_M_* that does not require the samples to be evenly distributed on the two PCs.

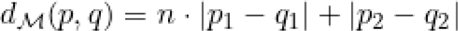

This distance matrix is formed by *d_M_* for each genomic location.

We also built performance evaluation scores for clustering quality and the number of genotypes present. The weight *n* in the above formula is chosen such that it generates the highest clustering quality. Furthermore, we calculate a membership probability that shows the probability of an individual belonging to each genotype cluster. The pipeline then produces a *gvcf* file that captures the information discussed above.

#### Identification of a novel QTL regulating short interspersed element (SINE) transposition rate

To demonstrate our ability to find short interspersed elements (SINEs), we assembled the genome of each strain using SuperNova, blasted the B2 SINE sequence against these 148 assemblies, and counted the number of high quality hits (e-score <1E-10, length > 149 nucleotides, identify > 75%). We then mapped B2 SINE counts using our marker map generated above.

#### Detection of BXD-private and shared variants

Methods for identifying high-confidence autosomal singleton variants are described in detail in a previous manuscript[31]. Briefly, we used *cyvcf2*[105] to identify single-nucleotide variants at which both founder genotypes (DBA/2J and C57BL/6J) and all but one of the BXD RILs were homozygous for the reference allele; at these sites, we therefore required a single BXD RIL (the focal line) to be either heterozygous or homozygous for the alternate allele. If the focal RIL was heterozygous for the alternate allele, we further required the fraction of reads supporting the alternate allele in that RIL to be >= 0.9. Finally, we required that the genotypes in both of the founders, as well as the focal RIL, were supported by >= 10 reads and had a Phred-scaled genotype quality of at least 20. We also removed all putative singletons that overlapped segmental duplications or simple repeat annotations in mm10/GRCm38, which were downloaded from the UCSC Genome Browser[106]. To identify epoch-private variants, we applied the same filters as previously described, but instead required the variant to be present in at least two of the sequenced BXDs, and for all of the BXDs with the shared variant to have the same parental haplotype. We also considered heterozygous genotypes with very high allele balance (i.e., the fraction of reads supporting the alternate allele >= 0.9) to be effectively homozygous. For candidate heterozygous and homozygous singletons (or candidate shared variants), we required the genotype call to be supported by at least 10 sequencing reads and have a Phred-scaled genotype quality of at least 20. Finally, we confirmed that at least one other BXD shared a parental haplotype identical-by-descent with the focal strain (i.e., the strain with the putative singleton) at the singleton site but was homozygous for the reference allele at that site.

We additionally annotated the full autosomal BXD VCF with SnpEff version 4.3t [107], using the GRCm38.86 database and the following command:

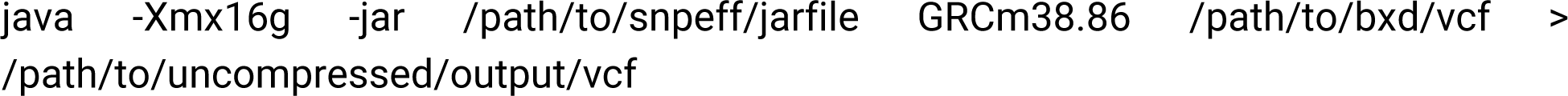

We estimated copy number states from read depth computed in non-overlapping 500bp sliding windows using *mosdepth*[108]. Specifically, the executed command was:

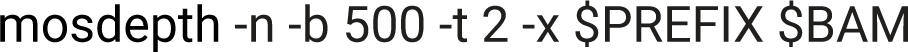

Raw read depth values were then corrected for potential GC-biases introduced during library preparation. Briefly, we used the mm10 reference genome to compute the observed GC content of each 500bp window. GC content values were rounded to the nearest 0.001 and regions with identical GC content were binned. For each strain, we then computed the mean read depth across all genomic windows that fell into each GC content bin. Next, we fitted a second degree polynomial to the relationship between read depth and GC content using the *scatter.smooth* function in R and with span parameter = 0.7. For each GC-bin, we then computed the difference between the fitted polynomial and the genome-wide average read depth. These values correspond to the magnitude of “inflation” or “deflation” in read depth across windows of a given GC-content due to systematic GC biases in the data. The read depth value in each 500bp window was then adjusted by the appropriate GC correction factor. Finally, these GC-corrected read depths were divided by the average per-sample coverage to convert into absolute copy number estimates. CN values across Chr X and Chr Y were then visualized using bedGraphs uploaded to IGV.

#### Genome-wide STR genotyping

We used Tandem Repeats Finder[109] (TRF) to identify regions within the mm10 mouse reference genome predicted to harbor STRs with repeat unit lengths up to 20bp using options matchscore=2; mismatchscore=5; indelscore=17; maxperiod=20; pm=80; pi=10; minscore=24; maxlen=1000. We processed this initial reference set using a custom script to (1) exclude homopolymer repeats which are highly error prone; (2) keep the shortest repeat unit for STRs that share either the same start or end position; (3) exclude “compound” repeats, consisting of multiple directly adjacent STRs with different repeat units; (4) trim repeat regions to only contain perfect repeats with no sequence imperfections; (5) collapse any duplicate STRs introduced by trimming; (6) remove overlapping STRs for which the repeat unit is identical; and (7) exclude very short repeats which we have observed are unlikely to be polymorphic. We filtered dinucleotide STRs with less than 5 perfect copies, trinucleotides with less than 4 perfect copies, and all other repeat classes with less than 3 perfect copies in mm10. After filtering, 1,176,016 STRs remained in our reference.

We used the “fix_read_length” development branch of GangSTR[37] (available at https://github.com/gymreklab/GangSTR/tree/fix_read_length) to genotype the reference STR loci in 152 BXD RI strains and the two founder strains C57BL/6J and DBA/2J from 10X Genomics Illumina short-read sequencing data. To reduce the effects of PCR “stutter” errors, we removed PCR duplicate reads using the –drop-dupes option. We then used HipSTR[110] to estimate per-locus stutter probabilities. We used a custom build to extend HipSTR to ignore the “AS/XS” BAM tags present in Chromium data, which are not properly currently handled in the HipSTR release, and to perform only stutter estimation but not genotyping. We ran HipSTR jointly on BAMs for all strains using our custom reference using option --min-reads 20 to output custom stutter models for each STR. For STRs at which HipSTR could not infer stutter models, including all repeats with repeat units >9bp, we set missing stutter parameters to default values of p=0.9; up=0.05; down=0.05. Additionally, since read length is a critical parameter used in GangSTR’s statistical model, we trimmed the second read in each read pair to 128bp to match the length of the first read. We then called GangSTR separately on each strain using our STR reference panel, trimmed and de-dupped reads, and per-locus stutter error probabilities as input. We additionally applied non-default parameter --max-proc-read, which was set to 4500 for DBA/2J which had higher coverage and 3000 for all other strains. This parameter skips loci with extremely high coverage which are likely to be error prone and consume high amounts of memory.

STR genotypes for each strain were filtered using the dumpSTR function from the TRTools package[111] with options `--min-call-DP 20; --max-call-DP 1000; --min-call-Q 0.9; --filter-badCI; --require-support 2; --readlen 128` to remove genotype calls with insufficient read depth, read support, or quality scores. Calls were then merged into a single multi-sample VCF file containing maximum likelihood diploid genotypes for each STR in each strain.

We applied the following filters to remove low-quality STRs from the merged VCF: (1) STRs overlapping known segmental duplication regions in the mm10 reference based on the mm10.genomicSuperDups table obtained from the UCSC Table Browser[106]; (2) STRs with call rates less than 90% across unfiltered strains; (3) STRs with no variation in repeat number across all strains; and (4) STRs for which variants from the mm10 reference were only observed in heterozygous genotypes. The final call-set contained 76,727 STR loci across 152 RI strains with an average per-strain call rate of 96.4%.

#### Validating STR genotypes using capillary electrophoresis

For each candidate STR, we designed primers to amplify the TR and surrounding region. A universal M13(-21) sequence (5’-TGTAAAACGACGGCCAGT-3’) was appended to each forward primer. We then amplified each TR using a three-primer reaction previously described^57^ consisting of the forward primer with the M13(-21) sequence, the reverse primer, and a third primer consisting of the M13(-21) sequence labeled with a fluorophore.

The forward (with M13(-21) sequence) and reverse primers for each TR were purchased through IDT. The labeled M13 primers were obtained through ThermoFisher (#450007) with fluorescent labels added to the 5’ ends (either FAM, VIC, NED, or PET). TRs were amplified using the forward and reverse primers plus an M13 primer with one of the four fluorophores with GoTaq polymerase (Promega #PRM7123) using PCR program: 94°C for 5 minutes, followed by 30 cycles of 94°C for 30 seconds, 58°C for 45 seconds, 72°C for 45 seconds, followed by 8 cycles of 94°C for 30 seconds, 53°C for 45 seconds, 72°C for 45 seconds, followed by 72°C for 30 minutes.

Fragment analysis of PCR products was performed on a ThermoFisher SeqStudio instrument using the GSLIZ1200 ladder, G5 (DS-33) dye set, and long fragment analysis options. Resulting .fsa files were analyzed using manual review in genemapper.

#### Long-read genome sequencing and data analysis

Two healthy adult male DBA/2J mice from the colony at the University of Tennessee Health Science Center were used. Spleen was used for DNA extraction. Oxford Nanopore (ONT) sequencing were conducted by using a Promethion instrument by DNA Link (Los Angeles, CA 90015). A total of 779,223 reads were obtained (14.25 billion bases, N50 of 29,205).

ONT data were mapped to the reference genome mm10 using minimap2 (version 2-2.17)[112]. SVs were detected using Sniffles (version 1.0.12)[107], SVIM (version 1.4.2)[113], and NanoVar (version 1.3.9)[114].

Pacific Biosciences (PacBio) HiFi data were generated by the DNA sequencing core facility at the University of Wisconsin. A total of 2,674,984 reads were obtained (28,568,875,629 bp, N50 of 11,307 and largest contig of 44,747). A total of 2,674,984 reads were obtained (28,568,875,629 bp, N50 of 11,307 and largest contig of 44,747).

#### Pangenome generation, and variant calling

Supernova haployd assemblies obtained by 10X linked reads were mapped against the GRCm38/mm10.fa reference genome using wfmash*v.0.6.0 (*https://github.com/ekg/wfmash) to select for reads mapping to chromosome 19. Mapped assemblies were used to build the pangenome with pggb *(v.0.2.0-pre+d8a5709-2)*[115] using the following combination of parameters:

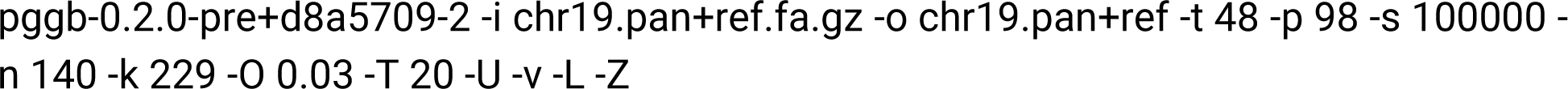

The variant calling on the pangenome was done using the following combination of parameters in vg*(v1.35.0-59-ge5be425c6)*[60].

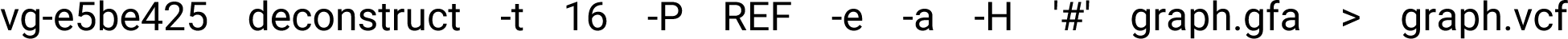

Validation of the call set was performed on DBA/2J using two true positive sets obtained from GATK *(v.4.0.3.0)* HaplotypeCaller on two sets of raw sequencing: the 10x and the PacBio sequence reads of DBA/2J. Prior to comparison the pangenome-derived, the PacBio-based, and the validated call sets were processed to remove missing data, sites where alleles are stretches of Ns, homozygous reference genotypes and variants greater than 50bp before normalization and decomposition using bcftools[116] under standard parameters. While the pangenome-derived VCF was based on haploid assemblies, for comparison purposes the calls were considered as homozygous diploid in the assumption that DBA/2J is fully isogenic, given ∼200 generations of sib-sib inbreeding. Comparison of the three call sets was performed with RTG tools *(v.3.12.1)*[117] using the *--squash-ploidy* option. RepeatMasker[118] was used to mask complex regions.

For pangenome graph visualization we used odgi[119] *(v.0.6.2)* and bandage[120] *(v. 0.8.1)*.

## Supplementary information

**Figure S1:**
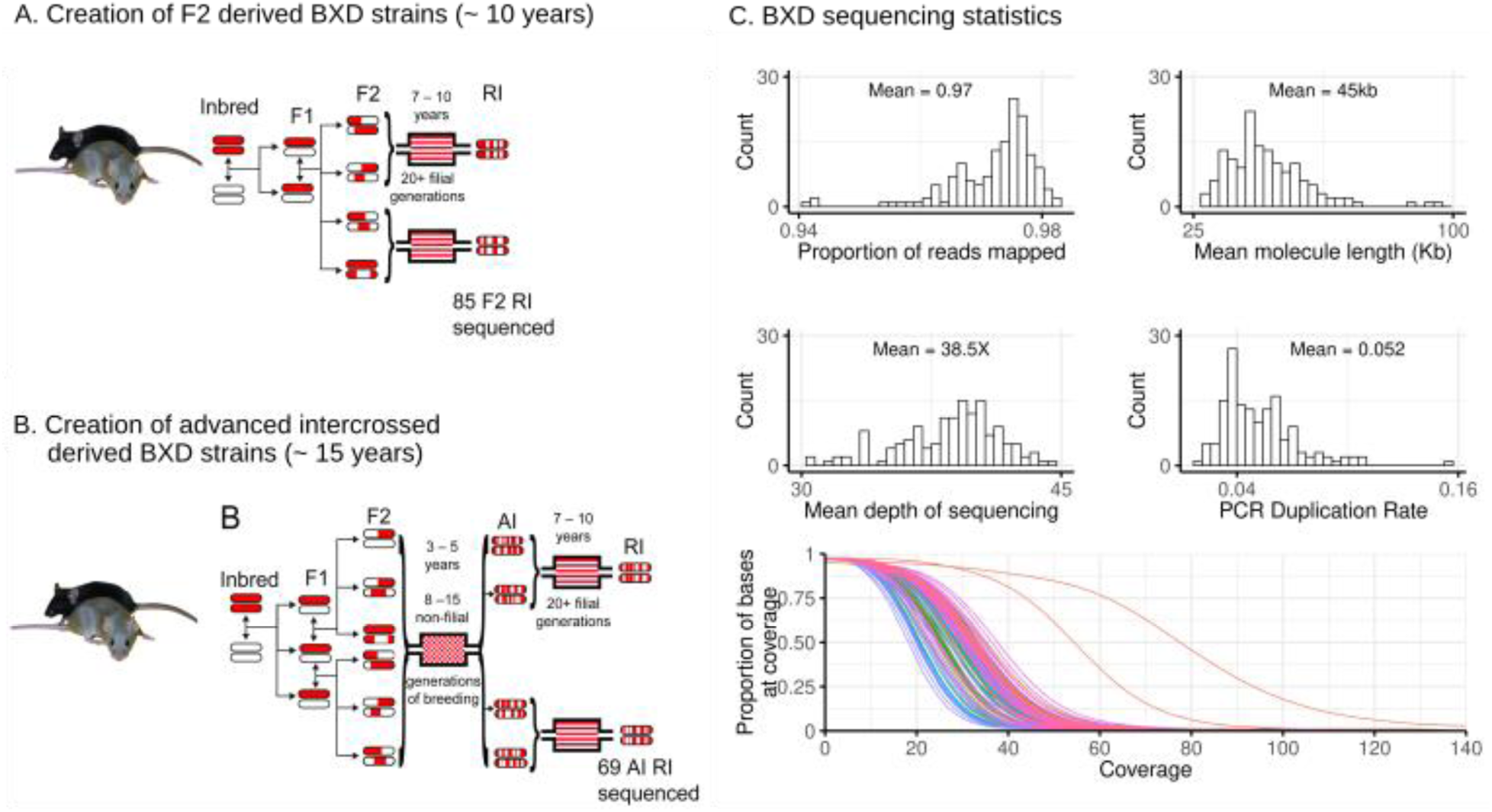
Creation of the BXD family (A, B), and sequencing statistics (C). The epochs of the BXD family were created using either a standard F2 (**A**, n = 85) or by advanced intercrosses (**B**, n = 69) from the C57BL/6J and DBA/2J parents in several epochs, from the 1970s, and the oldest of these strains have been bred for ∼200 generations **(Table S1)**. Red represents regions of the genome inherited from the C57BL/6J, and white represents regions of the genome coming from the DBA/2J. Solid lines represent a single generation. Adapted from[9,122]. This design results in both strain-specific variants, and variants between epochs. To detect these we used link read sequencing from 10X Genomics **(C)**, in which the genome is fragmented into ∼40Kb fragments, and these fragments are barcoded before further fragmentation and short read sequencing. Sequencing metrics across strains are given (**C**). Sequencing metrics across strains are given (**C**), showing high quality data with a mean proportion of mapped reads of 0.97, mean molecule length of 45kb, mean sequencing depth of 38.5X (after duplicate removal), and low PCR duplication rate of 5.2%. The coverage distribution curves (bottom panel) demonstrate the proportion of bases covered at different sequencing depths across all strains, with most strains showing similar coverage profiles except for two outlier samples with notably different coverage patterns.

**Figure S2:**
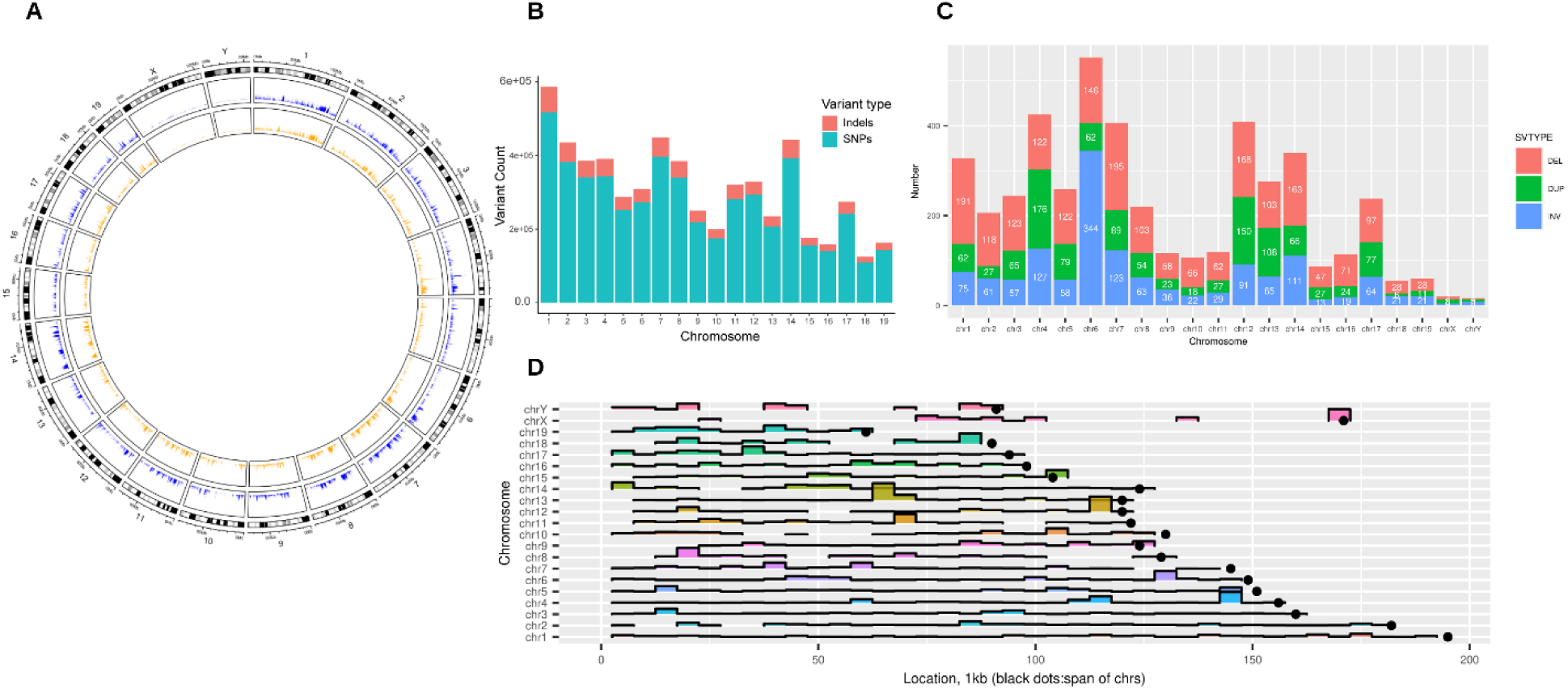
Quantitative results of small and large structural variant detection. **(A)** The distribution of SNPs (purple), Indels (orange), and STRs (green) across the BXD genome. Note areas that have a low number of all types of variants, indicating regions that are identical by decent between the C57BL/6J and DBA/2J strains. **(B)** Distribution of segregating (MAF > 0.2) SNPs and Indels across the autosomes of the BXD family **(C)** Number of structural variants (SVs). Deletions (red) are the most common form of SV detected. Variants are spread across the genome, although there is significant variation between chromosomes **(D)** Location of structural variants. There are SV hotspots throughout the genome.

**Figure S3:**
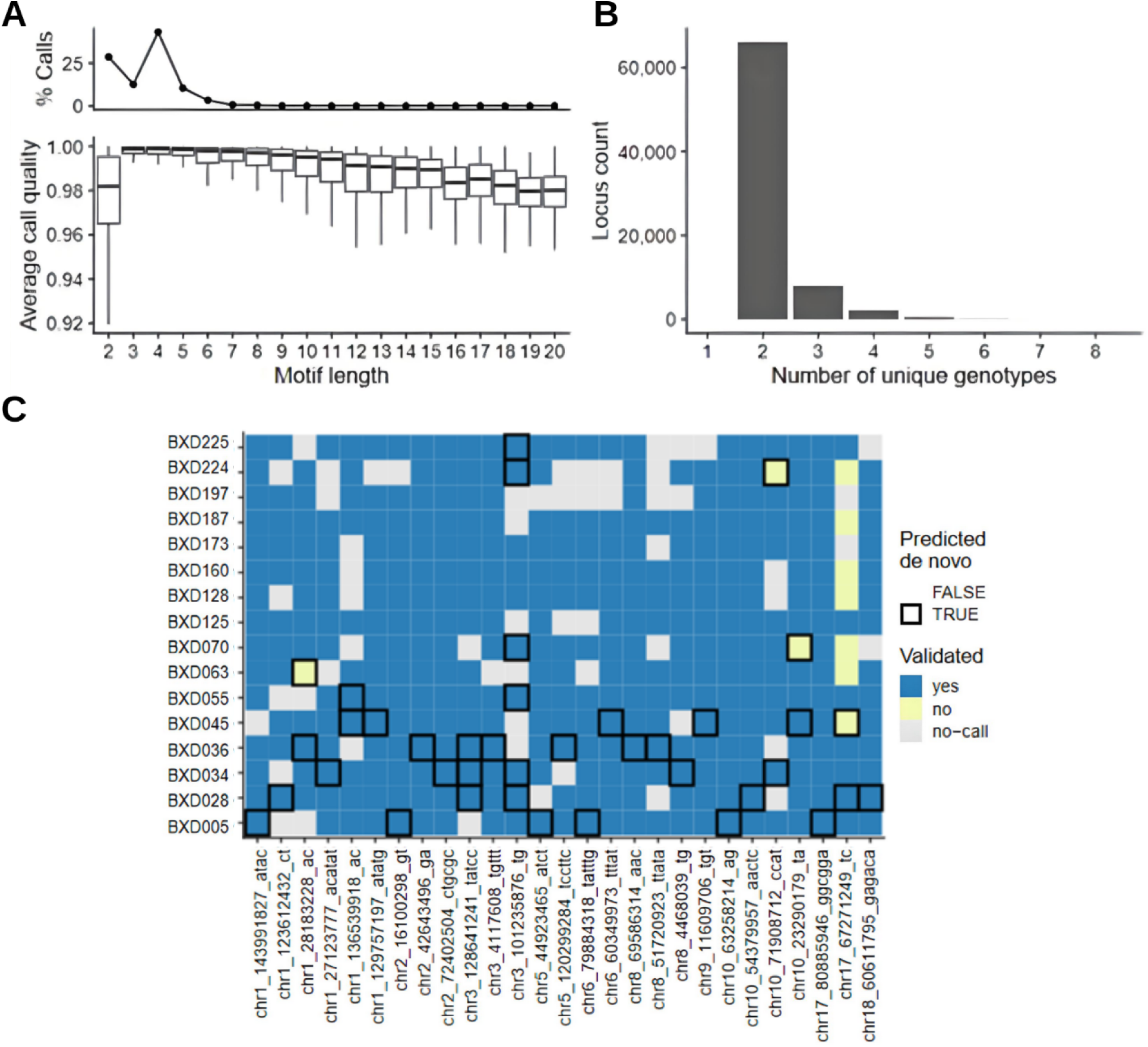
Summary of short tandem repeat (STR) identification. **(A)** proportion of (top panel) and average call quality of STRs of a given motif length in the dataset. **(B)** Distribution of the number of unique genotypes per locus for STRs. **(C)** STR genotype validation using capillary fragment analysis. Rows and columns represent strains and loci respectively, which were selected for validation. Heatmap fill indicates validation status: validated (blue), not validated (yellow). Strain/loci combinations missing a GangSTR call to compare to are filled in gray. Strain/loci combinations where a *de novo* STR variant was predicted by GangSTR are highlighted with a black border.

**Figure S4.**
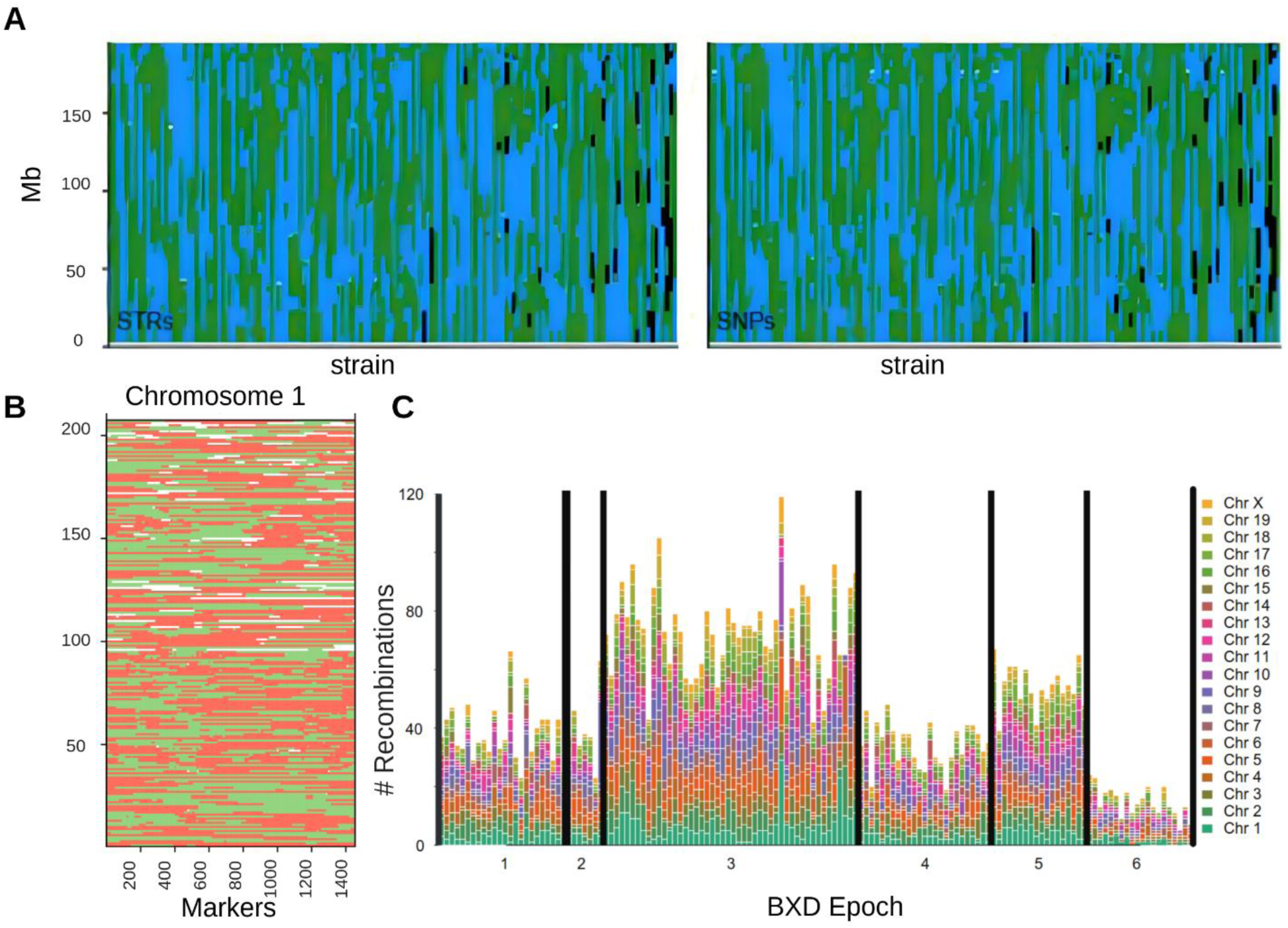
Recombinations per strain and precision of QTL mapping. **(A)** Homozygous patchwork of founder haplotype inheritance (C57BL/6J: green; DBA/2J: blue) for BXD RI strains, visualized as a strain by position matrix for STRs (left) and SNPs (right). **(B)** Recombination map of Chr 1. Regions of Chr 1 are shown as either C57BL/6J-like (red), DBA/2J-like (green) or unknown (white, either due to only having array-based genotypes available or because they were heterozygous at the time of sequencing). **(C)** The number of recombinations on each chromosome per strain, coloured by the chromosome, and divided by the epoch. It is clear to see that epochs 3 and 5, which were produced from advanced intercross lines, have more recombinations than those epochs produced from simple F2s.

## Supplementary Tables

Table S1 https://docs.google.com/spreadsheets/d/16WzQc1qM-ehDar8UPmVVQArr41QTI5i54aMVVsDm8Kg/edit?usp=sharing

**Table S2:**
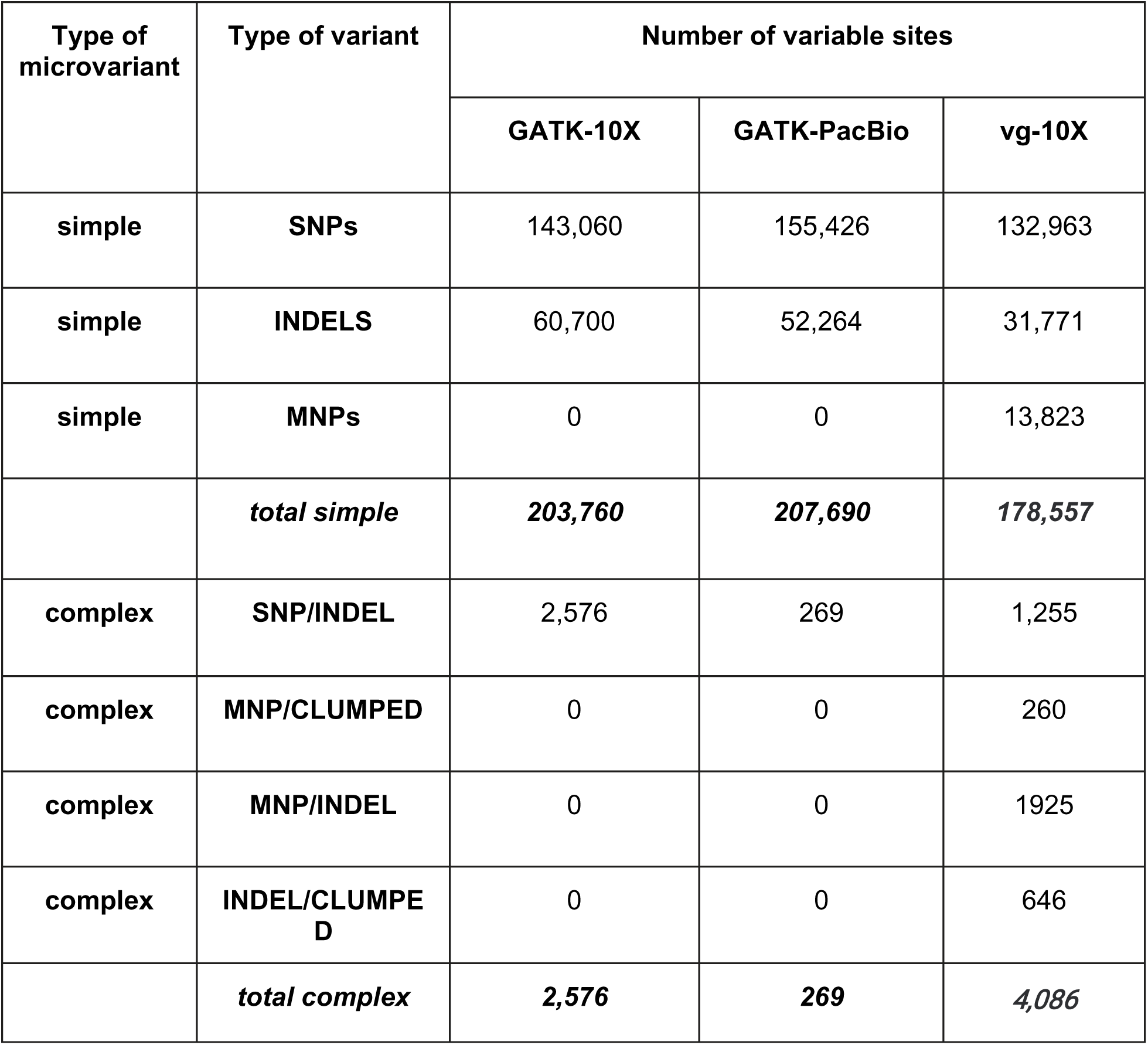
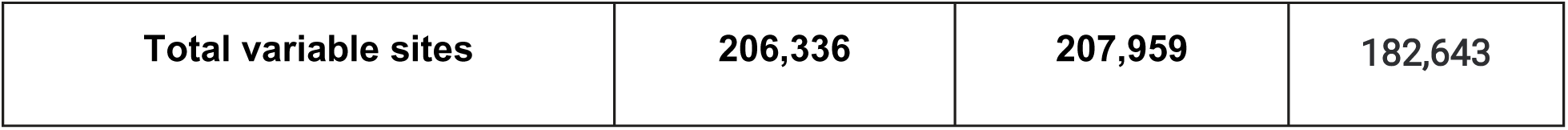
Variable sites on chromosome 19 in DBA/2J as called by GATK on genomics versus vg on pangenomic data. Truth sets are GATK on 10X (GATK-10X) and GATK on PacBio (GATK-PacBio) that were obtained by reference-based variant calling. Only microvariants (length up to 50bp) are considered.

**Table S3:** Quality of variant calling from the pangenome in DBA/2J evaluated on variants described in table S2. GK-10x: GATK on 10X sequence data; GK-PB: GATK on PacBio sequence data; vg-10X: vg on 10X sequence data. Gray columns (GK-10X vs GK-PB) are comparisons among the truth sets, in this case only the sequence technology is being evaluated.

|  |  | PRECISION (%) |  |  | SENSITIVITY (%) |  |  | F-MEASURE (%) |  |  |
| --- | --- | --- | --- | --- | --- | --- | --- | --- | --- | --- |
|  |  | GK-10X<br>vs<br>GK-PB | GK-10X<br>vs<br>vg-10X | GK-PB<br>vs<br>vg-10X | GK-10X<br>vs<br>GK-PB | GK-10X<br>vs<br>vg-10X | GK-PB<br>vs<br>vg-10X | GK-10X vs<br>GK-PB | GK-10X vs<br>vg-10X | GK-PB<br>vs<br>vg-10X |
| **Masked**<br>(37,677,383 bp) | **All** | 91.9 | 89.8 | 90.2 | 91.2 | 84.7 | 86.1 | 91.5 | 87.2 | 88.1 |
|  | **SNPs** | 95.3 | 93.2 | 94.1 | 97.3 | 85.3 | 84.3 | 96.3 | 89.1 | 88.9 |
|  | **Indels** | 78.3 | 88.5 | 86.3 | 67.7 | 51.8 | 57.4 | 73.8 | 65.3 | 69 |
| **Unmasked**<br>(23,742,621 bp) | **All** | 76.7 | 75.5 | 79.2 | 82.1 | 70.7 | 69.9 | 79.3 | 73 | 74.3 |

## Bibliography

1. Cubillos FA, Billi E, Zörgö E, Parts L, Fargier P, Omholt S, et al. Assessing the complex architecture of polygenic traits in diverged yeast populations. Mol Ecol. 2011;20:1401–13.

2. El-Din El-Assal S, Alonso-Blanco C, Peeters AJM, Raz V, Koornneef M. A QTL for flowering time in Arabidopsis reveals a novel allele of CRY2. Nat Genet. 2001;29:435–40.

3. Yin X. Model analysis of flowering phenology in recombinant inbred lines of barley. J Exp Bot. 2005;56:959–65.

4. Ruden DM, Chen L, Possidente D, Possidente B, Rasouli P, Wang L, et al. Genetical toxicogenomics in Drosophila identifies master-modulatory loci that are regulated by developmental exposure to lead. NeuroToxicology. 2009;30:898–914.

5. Cochrane BJ, Windelspecht M, Brandon S, Morrow M, Dryden L. Use of Recombinant Inbred Lines for the Investigation of Insecticide Resistance and Cross Resistance inDrosophila simulans. Pestic Biochem Physiol. 1998;61:95–114.

6. Snoek BL, Volkers RJM, Nijveen H, Petersen C, Dirksen P, Sterken MG, et al. A multi-parent recombinant inbred line population of C. elegans allows identification of novel QTLs for complex life history traits. BMC Biol. 2019;17:24.

7. Fitzgerald T, Brettell I, Leger A, Wolf N, Kusminski N, Monahan J, et al. The Medaka Inbred Kiyosu-Karlsruhe (MIKK) panel. Genome Biol. 2022;23:59.

8. Printz MP, Jirout M, Jaworski R, Alemayehu A, Kren V. Invited Review: HXB/BXH rat recombinant inbred strain platform: a newly enhanced tool for cardiovascular, behavioral, and developmental genetics and genomics. J Appl Physiol. 2003;94:2510–22.

9. Ashbrook DG, Arends D, Prins P, Mulligan MK, Roy S, Williams EG, et al. A platform for experimental precision medicine: The extended BXD mouse family. Cell Syst. 2021;12:235–247.e9.

10. Crow JF. Haldane, Bailey, Taylor and Recombinant-Inbred Lines. Genetics. 2007;176:729–32.

11. Taylor BA, Heiniger HJ, Meier H. Genetic Analysis of Resistance to Cadmium-Induced Testicular Damage in Mice. Exp Biol Med. 1973;143:629–33.

12. Taylor BA, Wnek C, Kotlus BS, Roemer N, MacTaggart T, Phillips SJ. Genotyping new BXD recombinant inbred mouse strains and comparison of BXD and consensus maps. Mamm Genome. 1999;10:335–48.

13. Sandoval-Sierra JV, Helbing AHB, Williams EG, Ashbrook DG, Roy S, Williams RW, et al. Body weight and high-fat diet are associated with epigenetic aging in female members of the BXD murine family. Aging Cell. 2020;19:e13207.

14. Wang X, Pandey AK, Mulligan MK, Williams EG, Mozhui K, Li Z, et al. Joint mouse–human phenome-wide association to test gene function and disease risk. Nat Commun. 2016;7:10464.

15. Roy S, Sleiman MB, Jha P, Ingels JF, Chapman CJ, McCarty MS, et al. Gene-by-environment modulation of lifespan and weight gain in the murine BXD family. Nat Metab. 2021;3:1217–27.

16. Wilkinson MD, Dumontier M, Aalbersberg IjJ, Appleton G, Axton M, Baak A, et al. The FAIR Guiding Principles for scientific data management and stewardship. Sci Data. 2016;3:160018.

17. Ashbrook DG, Mulligan MK, Williams RW. Post-genomic behavioral genetics: From revolution to routine. Genes Brain Behav. 2018;17:e12441.

18. Sloan Z, Arends D, W. Broman K, Centeno A, Furlotte N, Nijveen H, et al. GeneNetwork: framework for web-based genetics. J Open Source Softw. 2016;1:25.

19. Mulligan MK, Mozhui K, Prins P, Williams RW. GeneNetwork: A Toolbox for Systems Genetics. In: Schughart K, Williams RW, editors. Syst Genet [Internet]. New York, NY: Springer New York; 2017 [cited 2023 Oct 20]. p. 75–120. Available from: http://link.springer.com/10.1007/978-1-4939-6427-7_4

20. Mouse Genome Sequencing Consortium. Initial sequencing and comparative analysis of the mouse genome. Nature. 2002;420:520–62.

21. Church DM, Goodstadt L, Hillier LW, Zody MC, Goldstein S, She X, et al. Lineage-Specific Biology Revealed by a Finished Genome Assembly of the Mouse. Roberts RJ, editor. PLoS Biol. 2009;7:e1000112.

22. Bigot B. Use mouse biobanks or lose them.

23. Doran AG, Wong K, Flint J, Adams DJ, Hunter KW, Keane TM. Deep genome sequencing and variation analysis of 13 inbred mouse strains defines candidate phenotypic alleles, private variation and homozygous truncating mutations. Genome Biol. 2016;17:167.

24. Keane TM, Goodstadt L, Danecek P, White MA, Wong K, Yalcin B, et al. Mouse genomic variation and its effect on phenotypes and gene regulation. Nature. 2011;477:289–94.

25. Wang X, Agarwala R, Capra JA, Chen Z, Church DM, Ciobanu DC, et al. High-throughput sequencing of the DBA/2J mouse genome. BMC Bioinformatics. 2010;11:O7.

26. Li H, Auwerx J. Mouse Systems Genetics as a Prelude to Precision Medicine. Trends Genet. 2020;36:259–72.

27. Yalcin B, Adams DJ, Flint J, Keane TM. Next-generation sequencing of experimental mouse strains. Mamm Genome. 2012;23:490–8.

28. Lindsay SJ, Rahbari R, Kaplanis J, Keane T, Hurles ME. Similarities and differences in patterns of germline mutation between mice and humans. Nat Commun. 2019;10:4053.

29. Srivastava A, Morgan AP, Najarian ML, Sarsani VK, Sigmon JS, Shorter JR, et al. Genomes of the Mouse Collaborative Cross. Genetics. 2017;206:537–56.

30. Shorter JR, Najarian ML, Bell TA, Blanchard M, Ferris MT, Hock P, et al. Whole Genome Sequencing and Progress Toward Full Inbreeding of the Mouse Collaborative Cross Population. G3 GenesGenomesGenetics. 2019;9:1303–11.

31. Sasani TA, Ashbrook DG, Beichman AC, Lu L, Palmer AA, Williams RW, et al. A natural mutator allele shapes mutation spectrum variation in mice. Nature. 2022;605:497–502.

32. Maksimov MO, Wu C, Ashbrook DG, Villani F, Colonna V, Mousavi N, et al. A novel quantitative trait locus implicates *Msh3* in the propensity for genome-wide short tandem repeat expansions in mice. Genome Res. 2023;33:689–702.

33. Van Der Auwera GA, Carneiro MO, Hartl C, Poplin R, Del Angel G, Levy-Moonshine A, et al. From FastQ Data to High-Confidence Variant Calls: The Genome Analysis Toolkit Best Practices Pipeline. Curr Protoc Bioinforma [Internet]. 2013 [cited 2024 Dec 1];43. Available from: https://currentprotocols.onlinelibrary.wiley.com/doi/10.1002/0471250953.bi1110s43

34. DePristo MA, Banks E, Poplin R, Garimella KV, Maguire JR, Hartl C, et al. A framework for variation discovery and genotyping using next-generation DNA sequencing data. Nat Genet. 2011;43:491–8.

35. Gunturkun MH, Villani F, Colonna V, Ashbrook D, Williams RW, Chen H. SVJAM: Joint Analysis of Structural Variants Using Linked Read Sequencing Data [Internet]. Genomics; 2021 Nov. Available from: http://biorxiv.org/lookup/doi/10.1101/2021.11.02.467006

36. Marks P, Garcia S, Barrio AM, Belhocine K, Bernate J, Bharadwaj R, et al. Resolving the full spectrum of human genome variation using Linked-Reads. Genome Res. 2019;29:635–45.

37. Mousavi N, Shleizer-Burko S, Yanicky R, Gymrek M. Profiling the genome-wide landscape of tandem repeat expansions. Nucleic Acids Res. 2019;47:e90–e90.

38. Willems T, Zielinski D, Yuan J, Gordon A, Gymrek M, Erlich Y. Genome-wide profiling of heritable and de novo STR variations. Nat Methods. 2017;14:590–2.

39. Clarke LA. PCR amplification introduces errors into mononucleotide and dinucleotide repeat sequences. Mol Pathol. 2001;54:351–3.

40. Taft R, Davisson M, Wiles M. Know thy mouse. Trends Genet. 2006;22:649–53.

41. Sarsani VK, Raghupathy N, Fiddes IT, Armstrong J, Thibaud-Nissen F, Zinder O, et al. The genome of C57BL/6J “Eve”, the mother of the laboratory mouse genome reference strain [Internet]. 2019 [cited 2024 Dec 1]. Available from: http://biorxiv.org/lookup/doi/10.1101/517466

42. Mortazavi M, Ren Y, Saini S, Antaki D, Pierre CSt, Williams A, et al. Polymorphic SNPs, short tandem repeats and structural variants are responsible for differential gene expression across C57BL/6 and C57BL/10 substrains [Internet]. 2020 [cited 2024 Dec 1]. Available from: http://biorxiv.org/lookup/doi/10.1101/2020.03.16.993683

43. Watanabe T, Cheng E, Zhong M, Lin H. Retrotransposons and pseudogenes regulate mRNAs and lncRNAs via the piRNA pathway in the germline. Genome Res. 2015;25:368–80.

44. Rosen GD, Azoulay NG, Griffin EG, Newbury A, Koganti L, Fujisaki N, et al. Bilateral Subcortical Heterotopia with Partial Callosal Agenesis in a Mouse Mutant. Cereb Cortex. 2013;23:859–72.

45. Truong DT, Bonet A, Rendall AR, Rosen GD, Fitch RH. A Behavioral Evaluation of Sex Differences in a Mouse Model of Severe Neuronal Migration Disorder. Chapouthier G, editor. PLoS ONE. 2013;8:e73144.

46. Bi W, Sapir T, Shchelochkov OA, Zhang F, Withers MA, Hunter JV, et al. Increased LIS1 expression affects human and mouse brain development. Nat Genet. 2009;41:168–77.

47. Katayama K, Hayashi K, Inoue S, Sakaguchi K, Nakajima K. Enhanced expression of Pafah1b1 causes over-migration of cerebral cortical neurons into the marginal zone. Brain Struct Funct. 2017;222:4283–91.

48. Haverfield EV, Whited AJ, Petras KS, Dobyns WB, Das S. Intragenic deletions and duplications of the LIS1 and DCX genes: a major disease-causing mechanism in lissencephaly and subcortical band heterotopia. Eur J Hum Genet. 2009;17:911–8.

49. Brock S, Dobyns WB, Jansen A. PAFAH1B1-Related Lissencephaly / Subcortical Band Heterotopia.

50. Bryant CD, Smith DJ, Kantak KM, Nowak TS, Williams RW, Damaj MI, et al. Facilitating Complex Trait Analysis via Reduced Complexity Crosses. Trends Genet. 2020;36:549–62.

51. Eizenga JM, Novak AM, Sibbesen JA, Heumos S, Ghaffaari A, Hickey G, et al. Pangenome Graphs. Annu Rev Genomics Hum Genet. 2020;21:139–62.

52. Sirén J, Monlong J, Chang X, Novak AM, Eizenga JM, Markello C, et al. Pangenomics enables genotyping of known structural variants in 5202 diverse genomes. Science. 2021;374:abg8871.

53. Computational pan-genomics: status, promises and challenges. Brief Bioinform. 2016;bbw089.

54. Garrison E, Sirén J, Novak AM, Hickey G, Eizenga JM, Dawson ET, et al. Variation graph toolkit improves read mapping by representing genetic variation in the reference. Nat Biotechnol. 2018;36:875–9.

55. Wang F, Zhang Y, Yu X, Teng X-L, Ding R, Hu Z, et al. ZFP91 disturbs metabolic fitness and antitumor activity of tumor-infiltrating T cells. J Clin Invest. 2021;131:e144318.

56. Wang A, Ding L, Wu Z, Ding R, Teng X-L, Wang F, et al. ZFP91 is required for the maintenance of regulatory T cell homeostasis and function. J Exp Med. 2021;218:e20201217.

57. Villani F, Guarracino A, Ward RR, Green T, Emms M, Pravenec M, et al. Pangenome reconstruction in rats enhances genotype-phenotype mapping and variant discovery. 10.1016/j.isci.2025.111835

58. Théberge ET, Baker JA, Dubose C, Boyle JK, Balce K, Goldowitz D, et al. Genetic Influences on the Amount of Cell Death in the Neural Tube of BXD Mice Exposed to Acute Ethanol at Midgestation. Alcohol Clin Exp Res. 2019;43:439–52.

59. Zhou D, Zhao Y, Hook M, Zhao W, Starlard-Davenport A, Cook MN, et al. Ethanol’s Effect on Coq7 Expression in the Hippocampus of Mice. Front Genet. 2018;9:602.

60. Mulligan MK, Zhao W, Dickerson M, Arends D, Prins P, Cavigelli SA, et al. Genetic Contribution to Initial and Progressive Alcohol Intake Among Recombinant Inbred Strains of Mice. Front Genet. 2018;9:370.

61. Dickson PE, Roy TA, McNaughton KA, Wilcox TD, Kumar P, Chesler EJ. Systems genetics of sensation seeking. Genes Brain Behav. 2019;18:e12519.

62. Wang LS, Jiao Y, Huang Y, Liu XY, Gibson G, Bennett B, et al. Critical evaluation of transcription factor Atf2 as a candidate modulator of alcohol preference in mouse and human populations. Genet Mol Res. 2013;12:5992–6005.

63. Chella Krishnan K, Mukundan S, Alagarsamy J, Hur J, Nookala S, Siemens N, et al. Genetic Architecture of Group A Streptococcal Necrotizing Soft Tissue Infections in the Mouse. Bessen DE, editor. PLOS Pathog. 2016;12:e1005732.

64. Russo LM, Abdeltawab NF, O’Brien AD, Kotb M, Melton-Celsa AR. Mapping of genetic loci that modulate differential colonization by Escherichia coli O157:H7 TUV86-2 in advanced recombinant inbred BXD mice. BMC Genomics. 2015;16:947.

65. Boon AC, Williams RW, Sinasac DS, Webby RJ. A novel genetic locus linked to pro-inflammatory cytokines after virulent H5N1 virus infection in mice. BMC Genomics. 2014;15:1017.

66. Nedelko T, Kollmus H, Klawonn F, Spijker S, Lu L, Heßman M, et al. Distinct gene loci control the host response to influenza H1N1 virus infection in a time-dependent manner. BMC Genomics. 2012;13:411.

67. Boon ACM, deBeauchamp J, Hollmann A, Luke J, Kotb M, Rowe S, et al. Host Genetic Variation Affects Resistance to Infection with a Highly Pathogenic H5N1 Influenza A Virus in Mice. J Virol. 2009;83:10417–26.

68. Rodrigues BDA, Muñoz VR, Kuga GK, Gaspar RC, Nakandakari SCBR, Crisol BM, et al. Obesity Increases Mitogen-Activated Protein Kinase Phosphatase-3 Levels in the Hypothalamus of Mice. Front Cell Neurosci. 2017;11:313.

69. Jha P, McDevitt MT, Gupta R, Quiros PM, Williams EG, Gariani K, et al. Systems Analyses Reveal Physiological Roles and Genetic Regulators of Liver Lipid Species. Cell Syst. 2018;6:722–733.e6.

70. Jha P, McDevitt MT, Halilbasic E, Williams EG, Quiros PM, Gariani K, et al. Genetic Regulation of Plasma Lipid Species and Their Association with Metabolic Phenotypes. Cell Syst. 2018;6:709–721.e6.

71. Jones BC, Jellen LC. Systems Genetics Analysis of Iron and Its Regulation in Brain and Periphery. In: Schughart K, Williams RW, editors. Syst Genet [Internet]. New York, NY: Springer New York; 2017 [cited 2024 Dec 1]. p. 467–80. Available from: http://link.springer.com/10.1007/978-1-4939-6427-7_22

72. Reyes Fernandez PC, Replogle RA, Wang L, Zhang M, Fleet JC. Novel Genetic Loci Control Calcium Absorption and Femur Bone Mass as Well as Their Response to Low Calcium Intake in Male BXD Recombinant Inbred Mice. J Bone Miner Res. 2016;31:994–1002.

73. Fleet JC, Replogle RA, Reyes-Fernandez P, Wang L, Zhang M, Clinkenbeard EL, et al. Gene-by-Diet Interactions Affect Serum 1,25-Dihydroxyvitamin D Levels in Male BXD Recombinant Inbred Mice. Endocrinology. 2016;157:470–81.

74. Jung SH, Brownlow ML, Pellegrini M, Jankord R. Divergence in Morris Water Maze-Based Cognitive Performance under Chronic Stress Is Associated with the Hippocampal Whole Transcriptomic Modification in Mice. Front Mol Neurosci. 2017;10:275.

75. Diessler S, Jan M, Emmenegger Y, Guex N, Middleton B, Skene DJ, et al. A systems genetics resource and analysis of sleep regulation in the mouse. Kramer A, editor. PLOS Biol. 2018;16:e2005750.

76. King R, Lu L, Williams RW, Geisert EE. Transcriptome networks in the mouse retina: An exon level BXD RI database. Mol Vis. 2015;

77. Li H, Wang X, Rukina D, Huang Q, Lin T, Sorrentino V, et al. An Integrated Systems Genetics and Omics Toolkit to Probe Gene Function. Cell Syst. 2018;6:90–102.e4.

78. Parsons MJ, Grimm C, Paya-Cano JL, Fernandes C, Liu L, Philip VM, et al. Genetic variation in hippocampal microRNA expression differences in C57BL/6 J X DBA/2 J (BXD) recombinant inbred mouse strains. BMC Genomics. 2012;13:476.

79. Mulligan MK, DuBose C, Yue J, Miles MF, Lu L, Hamre KM. Expression, covariation, and genetic regulation of miRNA Biogenesis genes in brain supports their role in addiction, psychiatric disorders, and disease. Front Genet [Internet]. 2013 [cited 2024 Dec 1];4. Available from: http://journal.frontiersin.org/article/10.3389/fgene.2013.00126/abstract

80. Williams EG, Wu Y, Wolski W, Kim JY, Lan J, Hasan M, et al. Quantifying and Localizing the Mitochondrial Proteome Across Five Tissues in A Mouse Population. Mol Cell Proteomics. 2018;17:1766–77.

81. Williams EG, Pfister N, Roy S, Statzer C, Haverty J, Ingels J, et al. Multiomic profiling of the liver across diets and age in a diverse mouse population. Cell Syst. 2022;13:43–57.e6.

82. Williams EG, Wu Y, Jha P, Dubuis S, Blattmann P, Argmann CA, et al. Systems proteomics of liver mitochondria function. Science. 2016;352:aad0189.

83. Wu Y, Williams EG, Dubuis S, Mottis A, Jovaisaite V, Houten SM, et al. Multilayered Genetic and Omics Dissection of Mitochondrial Activity in a Mouse Reference Population. Cell. 2014;158:1415–30.

84. Baker CL, Walker M, Arat S, Ananda G, Petkova P, Powers NR, et al. Tissue-Specific Trans Regulation of the Mouse Epigenome.

85. Perez-Munoz ME, McKnite AM, Williams EG, Auwerx J, Williams RW, Peterson DA, et al. Diet modulates cecum bacterial diversity and physiological phenotypes across the BXD mouse genetic reference population. Wilson BA, editor. PLOS ONE. 2019;14:e0224100.

86. McKnite AM, Perez-Munoz ME, Lu L, Williams EG, Brewer S, Andreux PA, et al. Murine Gut Microbiota Is Defined by Host Genetics and Modulates Variation of Metabolic Traits. White BA, editor. PLoS ONE. 2012;7:e39191.

87. Li Z, Mulligan MK, Wang X, Miles MF, Lu L, Williams RW. A Transposon in Comt Generates mRNA Variants and Causes Widespread Expression and Behavioral Differences among Mice. Hoheisel J, editor. PLoS ONE. 2010;5:e12181.

88. Williams R, Lim JE, Harr B, Wing C, Walters R, Distler MG, et al. A Common and Unstable Copy Number Variant Is Associated with Differences in Glo1 Expression and Anxiety-Like Behavior. Heutink P, editor. PLoS ONE. 2009;4:e4649.

89. Chunduri A, Ashbrook DG. Old data and friends improve with age: Advancements with the updated tools of GeneNetwork [Internet]. 2021 [cited 2024 Dec 1]. Available from: http://biorxiv.org/lookup/doi/10.1101/2021.05.24.445383

90. Parker CC, Dickson PE, Philip VM, Thomas M, Chesler EJ. Systems Genetic Analysis in GeneNetwork.org. Curr Protoc Neurosci [Internet]. 2017 [cited 2024 Dec 1];79. Available from: https://currentprotocols.onlinelibrary.wiley.com/doi/10.1002/cpns.23

91. Watson PM, Ashbrook DG. GeneNetwork: a continuously updated tool for systems genetics analyses [Internet]. 2020 [cited 2024 Dec 1]. Available from: http://biorxiv.org/lookup/doi/10.1101/2020.12.23.424047

92. Collaborative Cross Consortium. The Genome Architecture of the Collaborative Cross Mouse Genetic Reference Population. Genetics. 2012;190:389–401.

93. The Complex Trait Consortium. The Collaborative Cross, a community resource for the genetic analysis of complex traits. Nat Genet. 2004;36:1133–7.

94. Threadgill DW, Churchill GA. Ten Years of the Collaborative Cross.

95. Chesler EJ, Miller DR, Branstetter LR, Galloway LD, Jackson BL, Philip VM, et al. The Collaborative Cross at Oak Ridge National Laboratory: developing a powerful resource for systems genetics. Mamm Genome. 2008;19:382–9.

96. The International Mouse Phenotyping Consortium, Dickinson ME, Flenniken AM, Ji X, Teboul L, Wong MD, et al. High-throughput discovery of novel developmental phenotypes. Nature. 2016;537:508–14.

97. A Mouse for All Reasons. Cell. 2007;128:9–13.

98. Collins FS, Finnell RH, Rossant J, Wurst W. A New Partner for the International Knockout Mouse Consortium. Cell. 2007;129:235.

99. Neuner SM, Garfinkel BP, Wilmott LA, Ignatowska-Jankowska BM, Citri A, Orly J, et al. Systems genetics identifies Hp1bp3 as a novel modulator of cognitive aging. Neurobiol Aging. 2016;46:58–67.

100. Neuner SM, Heuer SE, Huentelman MJ, O’Connell KMS, Kaczorowski CC. Harnessing Genetic Complexity to Enhance Translatability of Alzheimer’s Disease Mouse Models: A Path toward Precision Medicine. Neuron. 2019;101:399–411.e5.

101. O’Connell KMS, Ouellette AR, Neuner SM, Dunn AR, Kaczorowski CC. Genetic background modifies CNS-mediated sensorimotor decline in the AD-BXD mouse model of genetic diversity in Alzheimer’s disease. Genes Brain Behav. 2019;18:e12603.

102. Neuner SM, Heuer SE, Zhang J-G, Philip VM, Kaczorowski CC. Identification of Pre-symptomatic Gene Signatures That Predict Resilience to Cognitive Decline in the Genetically Diverse AD-BXD Model. Front Genet. 2019;10:35.

103. Simon SE, Simmons BW, Kim M, Joseph SC, Korba E, Marathe SJ, et al. Determining susceptibility loci in triple negative breast cancer using a novel pre-clinical model [Internet]. 2024 [cited 2024 Dec 1]. Available from: http://biorxiv.org/lookup/doi/10.1101/2024.02.08.579359

104. Cowin R-M, Bui N, Graham D, Green JR, Yuva-Paylor LA, Weiss A, et al. Genetic background modulates behavioral impairments in R6/2 mice and suggests a role for dominant genetic modifiers in Huntington’s disease pathogenesis. Mamm Genome. 2012;23:367–77.

105. Poplin R, Ruano-Rubio V, DePristo MA, Fennell TJ, Carneiro MO, Van Der Auwera GA, et al. Scaling accurate genetic variant discovery to tens of thousands of samples [Internet]. 2017 [cited 2024 Dec 1]. Available from: http://biorxiv.org/lookup/doi/10.1101/201178

106. Pedersen BS, Quinlan AR. cyvcf2: fast, flexible variant analysis with Python. Hancock J, editor. Bioinformatics. 2017;33:1867–9.

107. Navarro Gonzalez J, Zweig AS, Speir ML, Schmelter D, Rosenbloom KR, Raney BJ, et al. The UCSC Genome Browser database: 2021 update. Nucleic Acids Res. 2021;49:D1046–57.

108. Cingolani P, Platts A, Wang LL, Coon M, Nguyen T, Wang L, et al. A program for annotating and predicting the effects of single nucleotide polymorphisms, SnpEff: SNPs in the genome of Drosophila melanogaster strain w ^1118^ ; iso-2; iso-3. Fly (Austin). 2012;6:80–92.

109. Benson G. Tandem repeats finder: a program to analyze DNA sequences. Nucleic Acids Res. 1999;27:573–80.

110. Valle-Silva G, Frontanilla TS, Ayala J, Donadi EA, Simões AL, Castelli EC, et al. Analysis and comparison of the STR genotypes called with HipSTR, STRait Razor and toaSTR by using next generation sequencing data in a Brazilian population sample. Forensic Sci Int Genet. 2022;58:102676.

111. Mousavi N, Margoliash J, Pusarla N, Saini S, Yanicky R, Gymrek M. TRTools: a toolkit for genome-wide analysis of tandem repeats. Schwartz R, editor. Bioinformatics. 2021;37:731–3.

112. Li H. Minimap2: pairwise alignment for nucleotide sequences. Birol I, editor. Bioinformatics. 2018;34:3094–100.

113. Sedlazeck FJ, Rescheneder P, Smolka M, Fang H, Nattestad M, Von Haeseler A, et al. Accurate detection of complex structural variations using single-molecule sequencing. Nat Methods. 2018;15:461–8.

114. Heller D, Vingron M. SVIM: structural variant identification using mapped long reads. Birol I, editor. Bioinformatics. 2019;35:2907–15.

115. Tham CY, Tirado-Magallanes R, Goh Y, Fullwood MJ, Koh BTH, Wang W, et al. NanoVar: accurate characterization of patients’ genomic structural variants using low-depth nanopore sequencing. Genome Biol. 2020;21:56.

116. Garrison E, Guarracino A, Heumos S, Villani F, Bao Z, Tattini L, et al. Building pangenome graphs. Nat Methods. 2024;21:2008–12.

117. Danecek P, Bonfield JK, Liddle J, Marshall J, Ohan V, Pollard MO, et al. Twelve years of SAMtools and BCFtools. GigaScience. 2021;10:giab008.

118. Cleary JG, Braithwaite R, Gaastra K, Hilbush BS, Inglis S, Irvine SA, et al. Comparing Variant Call Files for Performance Benchmarking of Next-Generation Sequencing Variant Calling Pipelines [Internet]. Bioinformatics; 2015 Aug. Available from: http://biorxiv.org/lookup/doi/10.1101/023754

119. Tarailo-Graovac M, Chen N. Using RepeatMasker to identify repetitive elements in genomic sequences. Curr Protoc Bioinforma. 2009;Chapter 4:4.10.1-4.10.14.

120. Guarracino A, Heumos S, Nahnsen S, Prins P, Garrison E. ODGI: understanding pangenome graphs. Bioinforma Oxf Engl. 2022;38:3319–26.

121. Wick RR, Schultz MB, Zobel J, Holt KE. Bandage: interactive visualization of *de novo* genome assemblies. Bioinformatics. 2015;31:3350–2.

122. Peirce JL, Lu L, Gu J, Silver LM, Williams RW. A new set of BXD recombinant inbred lines from advanced intercross populations in mice. BMC Genet. 2004;5:7.

